# Inferring the Allelic Series at QTL in Multiparental Populations

**DOI:** 10.1101/2020.05.23.112326

**Authors:** Wesley L. Crouse, Samir N.P. Kelada, William Valdar

## Abstract

Multiparental populations (MPPs) are experimental populations in which the genome of every individual is a mosaic of known founder haplotypes. These populations are useful for detecting quantitative trait loci (QTL) because tests of association can leverage inferred founder haplotype descent. It is difficult, however, to determine how haplotypes at a locus group into distinct functional alleles, termed the allelic series. The allelic series is important because it provides information about the number of causal variants at a QTL and their combined effects. In this study, we introduce a fully-Bayesian model selection framework for inferring the allelic series. This framework accounts for sources of uncertainty found in typical MPPs, including the number and composition of functional alleles. Our prior distribution for the allelic series is based on the Chinese restaurant process, a relative of the Dirichlet process, and we leverage its connection to the coalescent to introduce additional prior information about haplotype relatedness via a phylogenetic tree. We evaluate our approach via simulation and apply it to QTL from two MPPs: the Collaborative Cross (CC) and the Drosophila Synthetic Population Resource (DSPR). We find that, although posterior inference of the exact allelic series is often uncertain, we are able to distinguish biallelic QTL from more complex multiallelic cases. Additionally, our allele-based approach improves haplotype effect estimation when the true number of functional alleles is small. Our method, Tree-Based Inference of Multiallelism via Bayesian Regression (TIMBR), provides new insight into the genetic architecture of QTL in MPPs.

---

Multiparental populations (MPPs) are experimental populations of model organisms generated by breeding a small but genetically diverse set of inbred parents to produce individual offspring whose genomes are mosaics of the original founder haplotypes (Churchill *et al.* 2004; Cavanagh *et al.* 2008; King *et al.* 2012). Because these haplotype mosaics succinctly describe the genetic differences between individuals, the standard approach for interrogating the genetic basis of quantitative traits in MPPs is haplotype-based association (Mott *et al.* 2000; Valdar *et al.* 2006; Aylor *et al.* 2011; Collaborative Cross Consortium *et al.* 2012; Huang *et al.* 2015; Broman *et al.* 2019).

The typical protocol for haplotype-based association is as follows. First, haplotypes are inferred along the genome for each individual by comparing its genotypes with those of the founders (Mott *et al.* 2000; Zheng *et al.* 2015; Broman *et al.* 2019). Then, quantitative trait locus (QTL) mapping proceeds by testing at each genomic locus the association of the trait with inferred founder haplotype state. For example, a linear model for the additive effect of *J* founder haplotypes at a genomic locus on a quantitative trait is given by

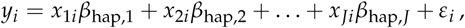

where *y*_*i*_ is the quantitative trait measurement of individual *i, x*_*ji*_ is the number of copies of the founder *j* haplotype, with *x*_*ji*_ ∈ {0, 1, 2} if haplotypes are known and *x*_*ji*_ ∈ [0, 2] if counts are estimated as imputed dosages, *β*_hap,*j*_ is the additive effect of the founder *j* haplotype, and *ε*_*i*_ is normally distributed error.

Haplotypes provide a richer source of information than observed variants (Haley and Knott 1992; Martínez and Curnow 1992). Whereas observed variant approaches such as single nucleotide polymorphism (SNP) association typically assume biallelic effects, haplotype-based association (technically a type of linkage disequilibrium mapping) tests the combined effects of all variants within the genomic interval, including any local epistastic interactions or variants that are unobserved or undiscovered (Zhang *et al.* 2014). This permits detection of complex genetic signals that may not be revealed by single-variant approaches, an advantage that has contributed to the widespread development of MPPs across a variety of biomedically (Churchill *et al*. 2004; Collaborative Cross Consortium *et al.* 2012; Macdonald and Long 2007; King *et al.* 2012; Kover *et al.* 2009) and agriculturally (Huang *et al.* 2015) important model organisms and species.

Nonetheless, the results of haplotype-based association do not translate directly into knowledge about causal variants. This is because the standard model used for haplotype-based association assumes that all haplotypes are functionally distinct and that their effects are independent. This latter assumption is biologically unlikely: it is more reasonable to expect that there are only a few causal variants at a locus, and that combinations of these variants will often be shared across haplotypes. More specifically, we expect that sets of shared causal variants partition the haplotypes into a potentially smaller number of functionally distinct alleles, with this assignment of haplotypes to functional alleles termed the allelic series.

An allelic series is easily superimposed onto the standard haplotype model by merging haplotypes into groups (Yalcin *et al.* 2005; Mosedale *et al.* 2019). For example, a linear model in the case of *J* = 5 haplotypes grouped into *K* = 3 functional alleles might be given by

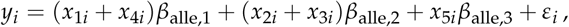

where haplotypes are functionally identical for founders 1 and 4, and for founders 2 and 3. In this case, the *J*-vector of haplotype effects ***β***_hap_ = (*β*_hap,1_, …, *β*_hap,*J*_)^*T*^ is collapsed into a *K*-vector of allele effects ***β***_alle_ = (*β*_alle,1_, …, *β*_alle,*K*_)^*T*^. This relationship is described in matrix notation by ***β***_hap_ = **M*β***_alle_, where **M** is a *J* × *K* indicator matrix that encodes the allelic series (Jannink and Wu 2003). Doing this, however, requires knowing the allelic series in advance, information that is not typically available.

Knowledge of the allelic series, and in particular, whether it is biallelic (*K* = 2) or multiallelic (*K* > 2), is critical for inference about the number of causal variants at a locus. This allelic perspective also suggests that the haplotype-based association approach is inefficient because it estimates redundant parameters when some haplotypes may be functionally equivalent (*K ≤ J*). Thus, an allele-based association approach would provide valuable insights into the number of causal variants while potentially improving effect estimation.

In this study, we introduce a method for QTL analysis that explicitly models an allelic series of haplotypes. Our method treats the allelic series as an unknown quantity that must be inferred from the data. In the context of the previous linear model, this means inferring the indicator matrix **M** while *K* is also unknown. This is a challenging problem because the number of possible allelic configurations is large even when the number of haplotypes is small.

There are currently no established methods for inferring the allelic series in MPPs, with QTL methods focused instead on, for example, accommodating uncertainty due to haplotype reconstruction (Mott *et al.* 2000; Kover *et al.* 2009; Durrant and Mott 2010; Zhang *et al.* 2014), or incorporating multiple QTL or terms for polygenic population structure (Valdar *et al.* 2009; Yuan *et al*. 2011; Gatti *et al.* 2014; Wei and Xu 2016). A recent study explored the relationship between the allelic series and QTL mapping power, but this was in the context of a haplotype-based association approach (Keele *et al.* 2019).

Inference of the allelic series in practice is often subjective, combining patterns in haplotype effect estimates with some intuition about the number of functional alleles (Aylor *et al.* 2011; Kelada *et al.* 2012). Yalcin *et al*. (2005) developed a method “merge analysis” that compares biallelic contrasts of “merged” haplotypes, as suggested by SNPs present in available sequence data, with the full haplotype model to see which most parsimoniously fits the data. This was trivially extended to multiallelic variants in Mosedale *et al*. (2019). In our framework, those approaches treat **M** and *K* as known and assume that they are implied by a single observed variant. King *et al*. (2014) also generalized merge analysis to interrogate the identity of the multiallelic contrast. Their approach implies a uniform prior distribution over the allelic series, p(**M**) ∝ 1. Their procedure, however, was ad hoc and not embedded within a broader statistical framework that could account for prior information about the allelic series.

The approach most closely resembling ours is Jannink and Wu (2003), used to infer the allelic series in doubled haploid lines. Their method places either a uniform or Poisson distribution on *K*, with the conditional allelic series then distributed uniformly, p(**M** |*K*) ∝ 1. They found that an allele-based model improves haplotype effect estimation but that inference of the allelic series itself was generally uncertain. Notably, their approach did not incorporate prior information about the relatedness of the haplotypes, which they identified as a key limitation. It is reasonable to expect that closely-related haplotypes are more likely to be functionally identical than distantly-related haplotypes, and consequently, that including this information would improve allelic series inference. Accounting for haplotype relatedness in an allele-based association framework is the primary innovation of our research.

Our approach frames inference of the allelic series as a Bayesian model selection problem. As suggested above, this requires specifying a prior distribution over the space of allelic configurations, p(**M**). Since this space is often much larger than the number of observations, information about the allelic series will typically be low. This makes the prior distribution critical, as it provides the basis for setting expectations about the number of functional alleles and their haplotype composition.

Our prior for p(**M**) is based on the Chinese restaurant process (CRP), which is the distribution over partitions that underlies the popular Dirichlet process mixture model (Escobar and West 1995; Müller *et al.* 2015). In this framework, the haplotypes are partitioned into a potentially smaller set of functional alleles, with the alleles having independent effects. The CRP allows for control over the prior number of alleles via its concentration parameter, but it implicitly assumes equal relatedness between individual haplotypes. We generalize the CRP prior to allow for unequal relatedness between the haplotypes by leveraging a particular property, namely, that the CRP can be described as the distribution of partitions induced by functional mutations on random coalescent trees, a representation known as Ewens’s sampling formula (Ewens 1972; Kingman 2006) (example in **Figure 1**).

**Figure 1.**
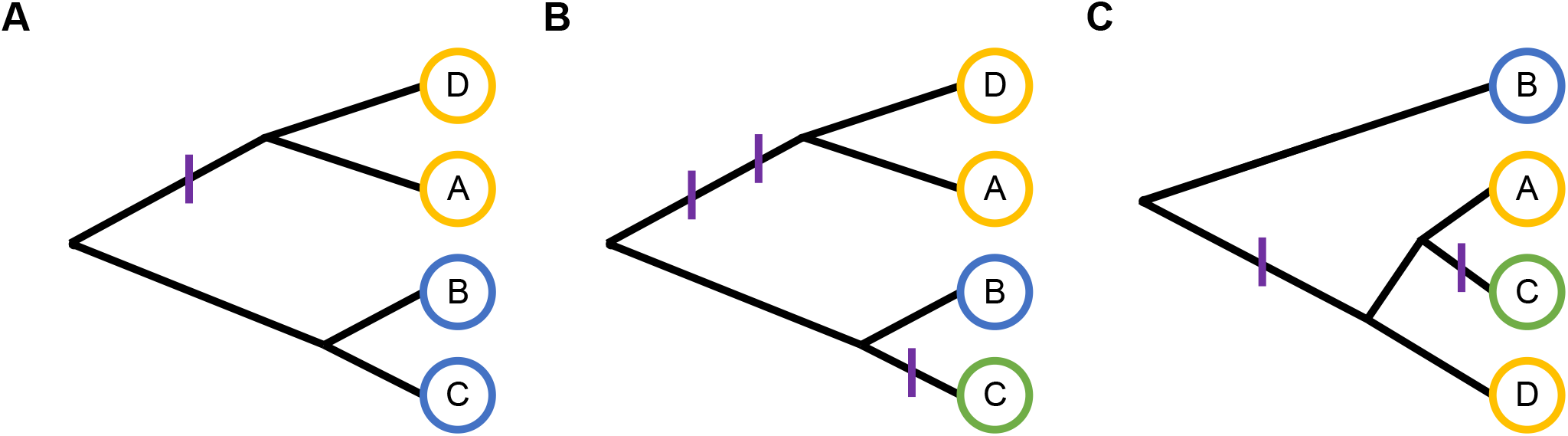
Allelic series induced by functional mutations on coalescent trees of haplotypes. Alleles are denoted by circle color (yellow, blue, green) and functional mutations are denoted by purple hashes. (**A**) One functional mutation on a tree partitions four haplotypes into two functional alleles: (A, D) | (B, C). (**B**) Additional mutations on the same tree partition the haplotypes into three functional alleles: (A, D) | (B) | (C). The second mutation does not affect (A, D). (**C**) Two functional mutations on a different tree partition the haplotypes into the same allelic series: (A, D) | (B) (C). Note that the allelic series from the first example, (A, D) | (B, C), is impossible given this tree.

Ewens’s sampling formula provides an intuitive mechanism for introducing prior information about haplotype relatedness: assuming that the phylogenetic tree of the haplotypes is known rather than random. This defines a prior distribution over the allelic series that is informed by a tree, p(**M**|*T*). In this way, our approach is similar to other models that include phylogenetic information; for example, by modeling distributional “change-points” on a tree (Azim Ansari and Didelot 2016), or by using phylogenetic distance as an input for a distance-dependent CRP (Cybis *et al.* 2018), among others (Zhang *et al.* 2012; Thompson and Kubatko 2013; Behr *et al.* 2020; Selle *et al.* 2020). In particular, Azim Ansari and Didelot (2016) specify a prior distribution over the allelic series by defining the prior probability that each branch of a tree is functionally mutated with respect to a phenotype (in their case, a categorical trait). This is also how we define p(**M**| *T*), and we highlight that this is embedded within the broader population genetics framework of Ewens’s sampling formula (Ewens 1972; Kingman 2006), with its connections to both the CRP and the coalescent (Berestycki 2009).

In the remainder, we introduce a fully-Bayesian framework for inferring the allelic series and additive allele effects in MPPs. This places the allelic series on a continuum that encompasses both single-variant and haplotype-based approaches at the limits. Our approach accounts for multiple sources of uncertainty found in typical MPPs, including uncertainty due to haplotype reconstruction, the number of functional alleles (Escobar and West 1995; Müller *et al.* 2015), and the magnitude of their effects (Gelman 2006). We outline a strategy for posterior inference using a partially-collapsed Gibbs sampler (Neal 2000; van Dyk and Park 2008; Park and Van Dyk 2009) and use posterior samples and Rao-Blackwellization to calculate the marginal likelihood (Blackwell 2007; Chib 1995), which is useful for comparing competing model assumptions (Kass and Raftery 1995). We then evaluate various properties of the allelic series approach via simulation and highlight several key findings. We conclude by presenting a series of illustrative real-data examples, from incipient lines of the Collaborative Cross (PreCC) (Kelada *et al*. 2012, 2014) and the Drosophila Synthetic Population Resource (DSPR) (King *et al.* 2014), that showcase the inferences facilitated by our allele-based approach.

## Materials and Methods

### Overview

At a quantitative trait locus (QTL), a trait **y** = (*y*_1_, …, *y*_*N*_)^*T*^ measured in *N* individuals *i* = 1, …, *N* is associated with genetic variation at a particular location in the genome. In a diploid multiparental population with *J ≥* 2 founder strains *j* = 1, …, *J*, this genetic variation is encoded by the pair of founder haplotypes, or the diplotype, at the locus, denoted for each individual by the indicator vector **d**_*i*_ with length 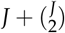. This length corresponds to the number of possible founder haplotype pairs, of which *J* are homozygous and 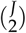 are heterozygous. The diplotype states of all individuals are given by the indicator matrix **D** = (**d**_1_, …, **d**_*N*_)^*T*^ with dimension 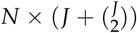. We are interested in understanding the the relationship between **y** and **D**.

For now, assume that the diplotype states are known, and that the phenotype is completely explained by the additive effects of the haplotypes and normally-distributed individual error, i.e. there are no other covariates, replicate observations, or population structure. Under these conditions, a linear model for **y** and **D** is

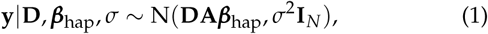

where ***β***_hap_ = (*β*_1_, …, *β* _*J*_)^*T*^ is a *J*-vector of haplotype effects, **A** is 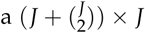 matrix that maps diplotypes state to haplotype frequency such that **DA** is an *N* × *J* design matrix of additive haplotype half-counts for each individual (i.e. the *ji*th element is half the number of founder *j* haplotypes at the locus for individual *i*), and *σ*^2^ scales the residual variance.

A standard Bayesian analysis of this linear model assumes that the haplotype effects are *a priori* distributed as

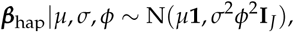

where *µ* is an intercept and *ϕ* controls the size of the haplotype effects relative to individual error (Servin and Stephens 2007). This fits an independent effect for each of the *J* founder haplotypes, implicitly assuming that, with respect to the phenotype, each founder haplotype is functionally distinct. This assumption, however, is rarely expected in practice. It is more realistic to assume that haplotypes group are grouped into *K ≤ J* functional alleles, with the assignment of haplotypes to functional alleles termed the allelic series.

Our approach extends the standard additive model to explicitly account for the allelic series, as in Jannink and Wu (2003). This decomposes the haplotype effects into the product of the allelic series matrix and a vector of allele effects:

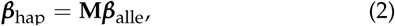

where ***β***_alle_ = (*β*_1_, …, *β*_*K*_)^*T*^ is a length *K* vector of allele effects, **M** = (**m**_1_, …, **m**_*J*_)^*T*^ is a *J* × *K* matrix denoting the allelic series, and **m**_*j*_ is a length *K* indicator vector denoting the allele assignment of strain *j*. For example, if there *J* = 3 haplotypes (labeled *A, B*, and *C*), and haplotypes *A* and *C* share one of *K* = 2 functional alleles, then the corresponding allelic series matrix is

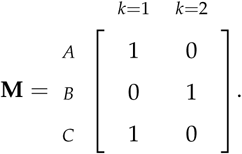

The haplotype effects ***β***_hap_ are not independent and functionally distinct, but instead are comprised of repeated values of a smaller set of allele effects ***β***_alle_. More generally, the allelic series matrix **M** partitions the *J* haplotypes into *K* functional alleles, which also determines the number of allele effects in ***β***_alle_. If the allelic series is known and *K* < *J*, this approach will estimate ***β***_hap_ more efficiently than the standard haplotype-based approach because it fits only *K* allele effects, rather than *J* redundant haplotype effects.

The allelic series is rarely known *a priori*, but it may be inferred from the data. From a Bayesian perspective, we are interested in the allelic series’ posterior distribution,

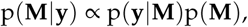

which requires specifying a prior distribution over the space of possible partitions encoded by **M**. Specifying a prior distribution over this space involves simultaneously defining expectations about the number of functional alleles and which combinations of haplotypes are more or less likely to be functionally distinct. This is particularly challenging when there are many founder haplotypes, as the space of allelic series partitions becomes exceedingly large.

Partition problems are common in Bayesian nonparametric statistics, and our approach is closely related to the popular Dirichlet process (DP) with a normal base distribution, i.e.,

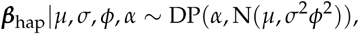

where *α* is the concentration parameter (Escobar and West 1995). Under the DP, the corresponding prior distributions on **M** and ***β***_alle_ are

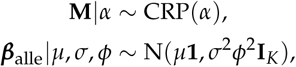

where CRP denotes the Chinese restaurant process. In the CRP, the concentration parameter *α* controls the prior distribution of the number of functional alleles, and the distribution over particular allelic configurations is implied by the process itself. Specifically, the CRP assigns a haplotype to an allele conditionally, in proportion to the number of haplotypes already assigned to that allele, without considering which particular haplotypes comprise the allele. In this way, the CRP is uninformative with respect to the relationship between individual haplotypes. We use the CRP as a starting point for directly modeling the allelic series, eventually modifying it in order to introduce additional prior information about haplotype relatedness.

The remainder of the methods first describes the likelihood function in more detail, using sum-to-zero contrasts and conjugate prior distributions to simplify the likelihood. Then, it focuses on the CRP, using Ewens’s sampling formula to show how it can be interpreted as a distribution over random coalescent trees with the haplotypes at the leaves. Next, this connection to the coalescent is used to define an informative prior distribution for the allelic series that reflects information about haplotype relatedness, as encoded by a phylogenetic tree. Prior distributions are then specified for the remaining model parameters, along with elicitation of prior hyperparameters. A graphical summary of the fully-specified model is shown in **Figure 2**. Next, we describe posterior inference via a partially-collapsed Gibbs sampler and how the output of this sampler can be used to estimate the marginal likelihood. The section ends by describing the simulations and real data examples presented in the results.

**Figure 2.**
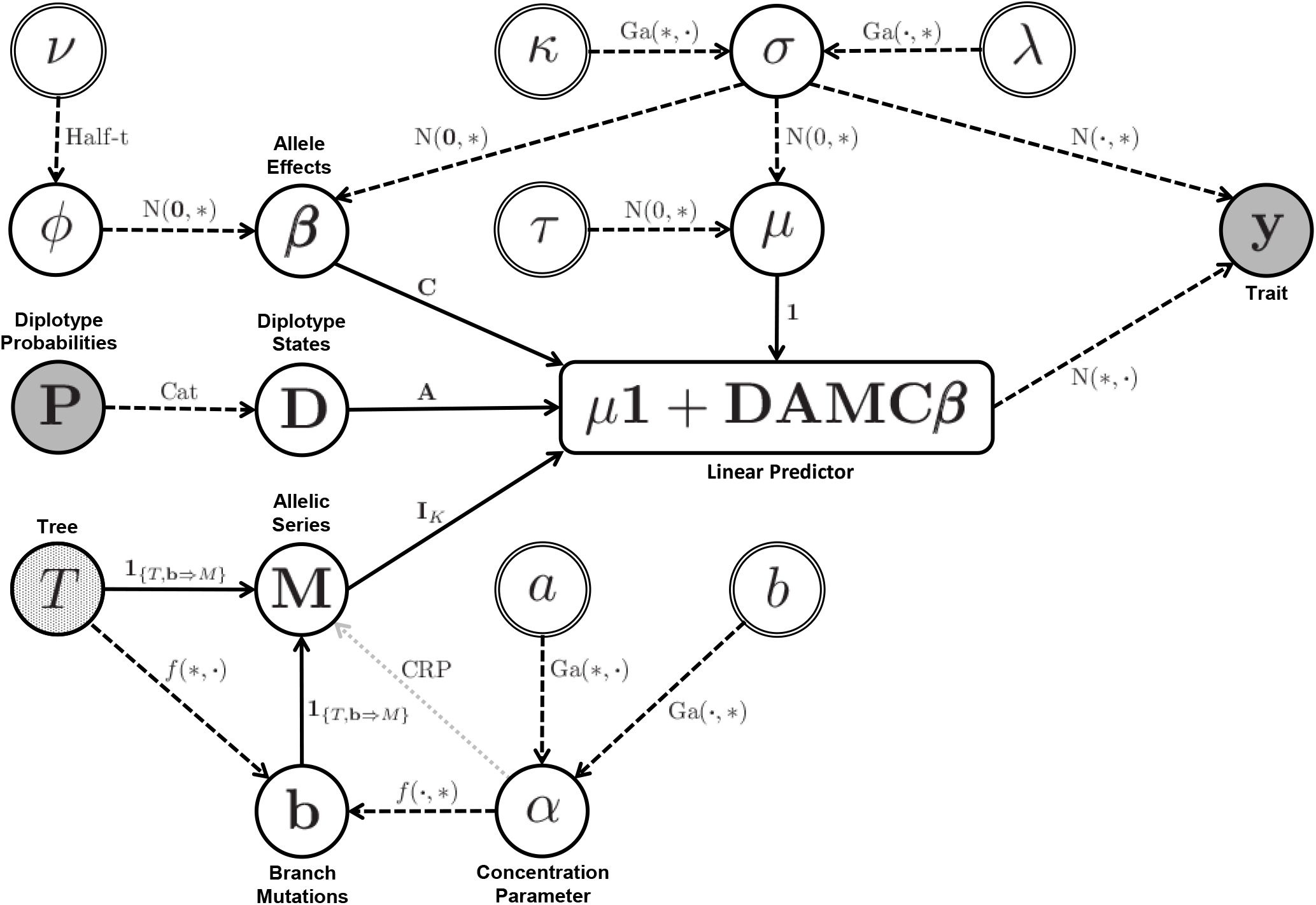
Graphical summary of the allele-based association approach. Shaded nodes are data, open nodes are variables, and double circle nodes are hyperparameters. Selected nodes are annotated to aid interpretation. Note that ***β*** is annotated as “allele effects” for brevity, but we actually depict the *K –* 1 independent effect vector, not ***β***_alle_. We also omit the subscripts on the hyperparameters *a*_*α*_ and *b*_*α*_. Arrows indicate dependencies between nodes. Dashed arrows are probabilistic dependencies, and labels denote the probability distributions linking the nodes. For distributions with multiple parameters, the star () indicates the parameter of the parent node, and the dot () is a placeholder for the other parameter. The notation *f* (*T, α*) is shorthand for the unnamed distribution p(**b**| *T, α*) given in the text. Solid arrows are deterministic dependencies, and labels denote the operation linking the nodes. The partial shading of the *T* node denotes that the tree can either be specified as data or treated as a variable with a coalescent prior distribution, which is parameterless. When *T* is a variable with a coalescent prior, *T* and *b* can be integrated from the model. Integration removes these nodes from the graph, leaving only the probabilistic dependency of **M** on *α*, given by the gray dotted arrow.

### Likelihood Function

The likelihood function defines the relationship between the phenotype and the diplotype states. Substituting **Equation 2** into **Equation 1** gives the likelihood function

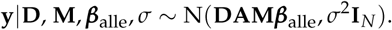

We make one additional substitution, reformulating the allele effects in terms of an intercept and a set of effects that are constrained to sum to zero,

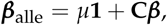

where **C** is a *K* × (*K –* 1) matrix of orthogonal sum-to-zero contrasts (Crowley *et al.* 2014) and ***β*** is a *K –* 1 vector of independent effects with mean zero. This substitution yields the likelihood

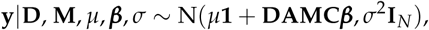

which now includes the intercept. In the expression **DAMC*β***, note that **A** and **C** are fixed, whereas **D, M**, and ***β*** are variables inferred by the model.

Our approach requires evaluating the likelihood over many settings of **M** with varying dimension. This motivates the use of conjugate priors for the allele effects, which allow us to simplify the likelihood by integrating, or collapsing, the allele effects out of the expression (Servin and Stephens 2007). Thus, we use the conjugate normal-gamma prior distribution for the precision, intercept, and independent effects:

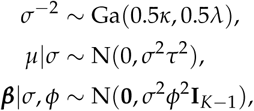

where *κ, λ* are shape and rate hyperparameters that control prior precision and *τ* is a hyperparameter that controls the informativeness of the prior intercept. Marginally, each allele effect is distributed according to

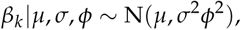

which is similar to the DP described earlier, but jointly, ***β***_*alle*_ is constrained to sum to *µ*.

The likelihood can be rewritten as

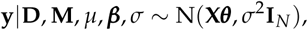

where **X** is the *N* × *K* design matrix of haplotype half-counts, **X** = [**1 DAMC]**, and ***θ***^*T*^ is a length *K* vector containing the intercept and independent effects, ***θ***^*T*^ = [*µ* ***β***^*T*^]. The intercept and independent effects are jointly distributed according to

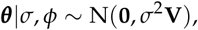

where **V** is a *K* × *K* diagonal matrix of the scaled prior covariance

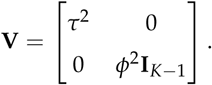

Conjugacy yields a closed form for a simplified, t-distributed likelihood function:

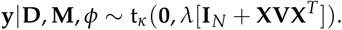

This simplified likelihood depends only on the diplotype states, allelic series configuration, and relative variance of the allele effects—this is useful during posterior inference. It is straight-forward to generalize the likelihood to account for covariates and replicate observations (**Appendix A**).

### Prior Distribution of the Allelic Series

#### Chinese Restaurant Process

Specifying a prior distribution over the allelic series involves defining expectations about the number of functional alleles and likely allelic configurations. This is challenging because the space of possible allelic series is large even when the number of haplotypes is small. For example, the Collaborative Cross (CC) has *J* = 8 founder haplotypes and 4,140 possible allelic series; the DSPR has *J* = 15 and over billion possibilities (Rota 1964). Encoding specific prior intuitions about such a space is difficult. It is tempting to consider a uniform prior over the allelic series [p(**M**) ∝ 1] that allows the likelihood to drive posterior inference about the allelic series. However, in most cases the number of observations will be much smaller than the number of possible allelic configurations, and this low-data scenario is precisely when prior information is most important. Instead of posterior inference being dominated by the likelihood, it will be subject to the properties of the uniform distribution, which include a strong prior belief in an intermediate number of functional alleles and a lack of flexibility to calibrate this belief.

Partition problems occur frequently in Bayesian nonparametric statistics, and a common and more flexible prior distribution is the CRP,

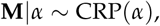

with probability density function

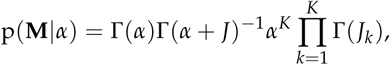

where *α* is a concentration parameter that controls the expected number of functional alleles, and *J*_*k*_ is the number of haplotypes assigned to allele *k* (Escobar and West 1995). The CRP is widely used in partition problems because it is exchangeable, making it amenable to posterior sampling. Exchangeability means that the density function of the CRP can be factored into conditional distributions that describe the allele assignment of a particular haplotype given the allelic configuration of all the other haplotypes. It also means that this conditional density can be applied iteratively (and in any order), beginning with all haplotypes unassigned, to construct the unconditional density of **M**|*α* (Welling 2006).

The conditional probability density function of the CRP is given by

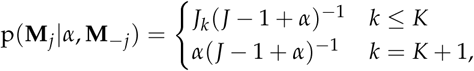

where **M**_*j*_ is the allele assignment of haplotype *j*, and **M** _−*j*_ is the allelic configuration of the other *J* − 1 haplotypes. The probability that haplotype *j* is assigned to allele *k* is proportional to the number of haplotypes already assigned to that allele, and the probability that haplotype *j* is assigned to a new allele is proportional to the concentration parameter *α*. This proportionality induces a “rich-get-richer” property that favors imbalanced allelic configurations (e.g. for *J* = 8, a biallelic contrast of 7 haplotypes vs 1 haplotype for *J* = 8, “7v1”) over balanced configurations (e.g. an even biallelic contrast, “4v4”) (Wallach *et al*. 2008). Note that the conditional probability that a haplotype is assigned to an existing functional allele does not depend on which particular haplotypes have already been assigned to that allele, only the number that have been assigned. In this way, the CRP is uninformative with respect to the relationship between individual haplotypes.

The CRP does, however, allow for control over the prior number of functional alleles via the concentration parameter. When *α* → ∞, all of the haplotypes will be assigned to a unique functional allele (**M** = **I**), which is identical to the standard haplotype approach that assumes that all J haplotypes are functionally distinct. When *α* → 0, all of the haplotypes will be assigned to a single functional allele (**M** = **1**), which is equivalent to a null model with no genetic effect.

To allow for additional flexibility, we place a prior distribution over the concentration parameter:

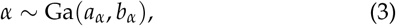

where *a*_*α*_ and *b*_*α*_ are hyperparameters that control the shape and rate of the concentration parameter. We discuss prior elicitation for these hyperparameters in a later subsection.

### Ewens’s Sampling Formula and the CRP

The CRP is equivalently given by Ewens’s sampling formula as the distribution over partitions induced by functional mutations on random coalescent trees with the founder haplotypes at the leaves (Ewens 1972; Kingman 2006). The intuition for this interpretation is as follows. At a QTL, there is a tree that describes the relatedness of the founder haplotypes. At various points during the evolution of this locus, functional mutations that altered the phenotype occurred at a constant rate on the branches of the tree. These functional mutations were transmitted to the founder haplotypes at the leaves of the tree, partitioning the haplotypes into groups that carry the same set of functional mutations. This partition is the allelic series. Examples of allelic series induced by functional mutations on coalescent trees of haplotypes are given in **Figure 1**. If we assume that the tree relating the founder haplotypes is unknown, but that it is distributed according to the coalescent process, then the resulting distribution over partitions is the CRP (Berestycki 2009).

More formally, Ewens’s sampling formula describes the allelic series as a function of a tree and which branches of that tree are functionally mutated. The conditional probability density function of the allelic series given a tree and branch mutations is

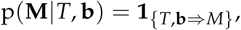

where *T* denotes a tree with *J* leaves and 2*J* − 2 branches, **b** = (*b*_1_, …, *b*_2*J*−2_)^*T*^ is a length 2*J* − 2 vector of indicators that denote if a branch is mutated, and **1** _{*T*,**b**_⇒_*M*}_ is an indicator function that takes value 1 when *T* and **b** imply **M** and 0 otherwise.

The tree *T* is an unknown random graph that is distributed according to the coalescent process with *J* leaves:

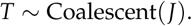

Coalescent trees are defined by sequential coalescent events that join lineages of the tree in random order, beginning with the leaves, as well as the times at which these coalescent events occur, which are exponentially distributed and depend on the number of lineages remaining prior to each coalescence (Kingman 1982). For our purposes, it is sufficient to note that there is a probability distribution over trees, p(*T*), and that this distribution assumes equal relatedness of the haplotypes via the random order of coalescent events. We also note that each branch of the tree has a corresponding length, which is contained in the length 2*J –* 2 vector 𝓁 and described in coalescent units.

The mutation status of the branches **b** is an unknown vector of indicators. Assume that functional mutations occur on the branches of the tree as a Poisson process with constant rate 0.5*α*. Then the number of mutations on each branch is Poisson distributed with rate proportional to branch length, and the probability density function for **b**, which indicates whether or not each branch is mutated is

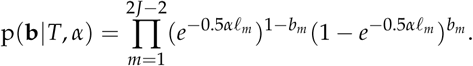

This is similar to Azim Ansari and Didelot (2016), but with branch lengths scaled by 0.5 and in coalescent units. The concentration parameter *α* controls the functional mutation rate per half-unit of coalescent branch length. Note that moving forward, we will refer to *α* interchangeably as the concentration parameter (of the CRP) or the functional mutation rate (on the tree), depending on context. When *α* → ∞, the probability that each branch is mutated approaches 1, the tree is saturated with functional mutations, and all of the founder haplotypes are functionally distinct (**M** = **I**). When *α* → 0, the probability that each branch is mutated approaches 0, there are no functional mutations on the tree, and all of the founders are functionally identical (**M** = **1**).

The probability density function for the allelic series is thus

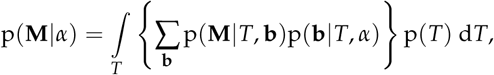

which involves identifying the allelic series implied by each combination of mutated branches on a tree, weighing by the probability of that combination, summing over all possible combinations, and then integrating over all possible coalescent trees. Remarkably, this is identical to the probability density function of the CRP described previously (Berestycki 2009). In this framework, the “rich-get-richer” property of the CRP is induced by integrating over coalescent tree structures (e.g. all trees with J=8 haplotypes have a branch that permits a specific 7v1 contrast, whereas only a subset of these trees have a branch that permits a specific 4v4 contrast; see **Figure 1** for an illustration).

#### Tree-Informed CRP

The CRP described above permits prior control of the number of functional alleles but, by integrating over all possible coalescent trees from random coalescent lineages, assumes that haplotypes are equally related. Specifying unequal relatedness in this framework is straightforward, however, if haplotype relationships can be specified in a tree. Conditional on a tree, the distribution over the allelic series reflects the relationships defined by the structure of the tree and the lengths of its branches. The tree topology reduces the space of possible partitions because many settings of **M** violate the relationships defined by *T*, making this information highly informative. The branch lengths of *T* also provide information about the allelic series, as long branches are more likely to be functionally mutated than short branches. Consequently, haplotypes separated by longer branches are more likely to be functionally distinct than haplotypes separated by shorter branches. The functional mutation rate still controls the prior number of functional alleles, now in combination with the tree structure and branch lengths.

If the tree is known, the conditional probability density function of the allelic series is given by

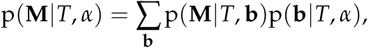

which does not involve integrating over coalescent trees, but does involve (weighted) summation over all 2^2*J*−2^ possible configurations of **b**. This approach is computationally intractable when the number of haplotypes is large, but provided *J* is small (e.g. *J* = 8, the case for many MPPs, but not *J* = 15, the case for the DSPR), it is possible to compute p(**M** |*T, α*) directly. We focus on this approach and consider alternatives in the discussion.

Recall that the functional mutation rate (concentration parameter) *α* is an unknown variable with a prior distribution. To avoid calculating p(**M**|*T, α*) for many settings of *α* during posterior inference, we marginalize over this variable and compute p(**M**|*T*) directly. The conditional probability density function is given by

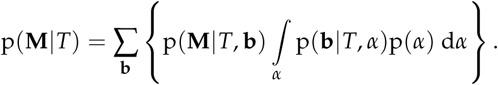

In **Appendix B**, we show that the integral over *α* can be computed exactly when *α* has a gamma prior distribution.

Lastly, to this point, we have assumed that the tree is known, but it may be unknown and inferred with uncertainty from a sequence alignment (Drummond *et al.* 2012). In this case, we are interested in the allelic series prior distribution conditional on the sequence alignment *S*,

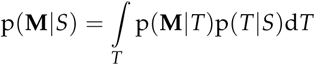

This can be approximated by averaging p(**M**|*T*) over a sample of trees from p(*T*|*S*).

### Prior Distribution of Diplotype States

The diplotype state of each individual is an unobserved latent variable that is probabilistically inferred via haplotype reconstruction. To account for this uncertainty, the diplotype state of each individual is given a categorical prior distribution

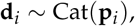

where **p**_*i*_ is a 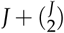 length vector of prior diplotype probabilities for each individual.

### Prior Distribution of Effect Size

The variable *ϕ* controls the size of the allele effects relative to individual error and controls the degree to which model complexity is penalized in Bayesian regression. We place a half-t prior distribution on the scaled standard deviation of the allele effects:

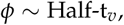

where *v* is degrees of freedom of the half-t distribution. The half-t is a preferred prior choice for variance components in normally-distributed models due to its behavior at the boundary (*ϕ* = 0) and its convenient representation as the product of two conditionally-conjugate latent variables (Gelman 2006).

### Prior Elicitation

This subsection describes the selection of hyperparameters for the priors describe above and discusses relevant considerations that may influence this selection.

#### Individual Error and Intercept

We use uninformative prior distributions for the size of the individual error and the intercept of the data. Specifically, we assume the limiting form of the prior distributions for *σ* as *κ, ϕ* → 0 and *µ* as *τ* → ∞. These prior distributions are improper, but posterior inference is still proper when these quantities are informed by the data (Servin and Stephens 2007).

#### Concentration Parameter / Functional Mutation Rate

The shape and rate parameters *a*_*α*_ and *b*_*α*_ control the prior distribution of the concentration parameter (functional mutation rate), which in turn controls the prior distribution over the number of functional alleles. An uninformative prior distribution for the concentration parameter is given by *a*_*α*_, *b*_*α*_ → 0 (Escobar and West 1995). Posterior learning about the concentration parameter, however, depends only on the number of founder haplotypes *J* and the number of functional alleles *K*. For this reason, even if **M** is known, the concentration parameter is poorly informed when *J* is small. This necessitates a prior distribution that reflects reasonable prior expectations about the number of functional alleles.

We focus on one particular prior distribution for the concentration parameter: an exponential distribution that places 50% of the prior probability on the null model, given by *a*_*α*_ = 1 and *b*_*α*_ ≈ 2.33 when *J* = 8. This prior distribution favors small numbers of functional alleles with low variance. In the Supplement, we consider two alternative prior distributions that favor higher numbers of alleles with varying degrees of certainty. There are many ways to calibrate prior expectations about the number of alleles, for example, by considering the frequency of biallelic contrasts, or the expected number of functional mutations on a tree, as in Azim Ansari and Didelot (2016). We emphasize that the reasonableness of a prior is specific to the number of founder haplotypes, the nature of the analysis (pre- or post-QTL detection), and other population- or trait-specific prior beliefs.

#### Coalescent Tree

Specifying a prior tree for the haplotypes is highly informative with respect to the allelic series. Our framework assumes that the phylogenetic tree is coalescent (with branches in coalescent units), satisfying assumptions of no recombination, selection or population structure. In the context of QTL mapping, the exact location of the causal sequence is often uncertain, making it difficult to satisfy the assumption of no recombination in particular. We discuss inferring trees in recombinant organisms in more detail in the discussion section, and we evaluate the consequences of tree misspecification in our simulations. By default, we recommend using the CRP, which assumes an unknown coalescent tree.

#### Diplotype States

It is assumed that the prior diplotype state probabilities of each individual **p**_*i*_ have been previously inferred from genotype or sequence data using established methods for haplotype reconstruction (Mott *et al.* 2000; Zheng *et al.* 2015; Broman *et al.* 2019).

#### Relative Allele Effect Size

The half-t prior distribution on *ϕ*, the scaled standard deviation of allele effect size, is controlled by degrees of freedom *v*. We set *v* = 2, which is the minimum value of *v* that yields a monotonically decreasing prior distribution for the proportion of variance explained by the QTL, given by 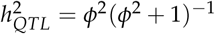. Larger values of *v* reflect a stronger prior belief in “small” effect sizes and increase the degree of shrinkage in the allele effects.

### Posterior Inference

Posterior samples are obtained using a partially-collapsed Gibbs sampler (van Dyk and Park 2008; Park and Van Dyk 2009). This involves four general steps:

1. updating the allelic series with the effects and scale of the error integrated, or collapsed, from the model,
2. jointly sampling the error scale and effects,
3. updating the relative size of the allelic effects, and
4. jointly updating the diplotype states.

The effects and scale of the error are integrated from the model during the first step in order to avoid mismatching the dimension of ***β*** and the dimension of **M** when updating the allelic series. After updating the allelic series, the effects and scale of the error are reintroduced into the model in order to take advantage of a convenient latent variable sampling scheme for the relative size of the allele effects and to facilitate a joint update of the diplotype states. These steps are discussed in more detail below.

#### 1a. Updating the Allelic Series with the CRP

We update the allele assignment of each haplotype individually, conditional on the allele assignment of the other haplotypes. In the case of the CRP, we must also update the concentration parameter. The conditional posteriors of the allelic series and the concentration parameter under the CRP are given by

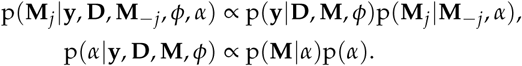

The first equation is the product of the t-distributed likelihood and the categorical, exchangeable, conditional prior distribution of the CRP. The conditional posterior is calculated directly by evaluating the likelihood at all possible (conditional) settings of the allelic series (Neal 2000). The conditional posterior of the concentration parameter depends only on the number of alleles in the allelic series, and there is a convenient, well-established latent variable approach for sampling from this posterior distribution (Escobar and West 1995; Müller *et al.* 2015).

#### 1b. Updating the Allelic Series with a Tree

In the case of the tree-informed prior distribution, the concentration parameter has already been integrated from the allelic series prior. Thus, the conditional posterior of the allelic series under the tree-informed prior is given by

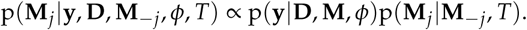

which is similar to the previous equation. The conditional prior distribution of the allelic series given the tree, p(**M**_*j*_ | **M**_**–***j*_, *T*), however, is not exchangeable and is not easy to calculate. Thus, we assume that the conditional prior distribution is proportional to the marginal prior distribution

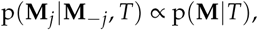

treating this distribution as if it were exchangeable. At each iteration of the sampler, we randomize the order in which the haplotypes assignments are updated. This is to avoid bias introduced by ordered updates of nonexchangeable variables, as described by Wallach *et al*. (2008) in the context of a uniform process. We have not observed issues with mixing using this approach, suggesting that this violation of exchangeability is mild. We outline an alternative, exchangeable approach for this step in **Appendix C**

#### 2. Sampling the Error Scale and Effects

The conditional posterior of the intercept, allele effects, and error scale is given by

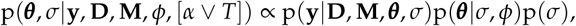

which is the product of the normally-distributed likelihood and a conjugate normal-gamma prior distribution, yielding a normal-gamma conditional posterior distribution, as in Servin and Stephens (2007). The notation [*α* ∨ *T*] denotes that the distribution is conditional on either *α* or *T*, depending on which allelic series prior distribution is used.

#### 3. Updating the Relative Size of the Allele Effects

The conditional posterior of the relative size of the allele effects is given by

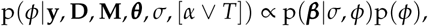

which is the product of a normal distribution and a half-t prior distribution for the scaled standard deviation, which is not conjugate. The half-t prior distribution, however, can be re-expressed as the product of two latent variables: the square root of an inverse-gamma-distributed variable and the absolute value of a normally-distributed variable. Respectively, these variables are conditionally conjugate to the prior distribution of ***β*** and the likelihood function, allowing for straightforward sampling of these latent variables (Gelman 2006).

#### 4. Updating the Diplotype States

The conditional posterior of the diplotype states is given by

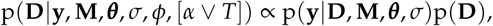

which is the product of the normally-distributed likelihood and categorical prior distributions. The diplotype states are conditionally independent of each other, and the joint conditional posterior is computed by evaluating the likelihood of each individual observation over all possible diplotype states.

### Marginal Likelihood

The marginal likelihood is useful for evaluating different hypotheses about the data. For example, two competing hypothesis can be evaluated by calculating their Bayes factor (BF), which is the ratio of the marginal likelihoods under each hypothesis (Kass and Raftery 1995). In **Appendix D**, we outline an approach for estimating the marginal likelihood using the output of the Gibbs sampler (Chib 1995). We use this estimate to compute BFs in favor of the allele-based approach versus the haplotype-based approach. There is precedent for the use of BFs in statistical genetics (Servin and Stephens 2007), but we note that BFs are not without criticism (Robert 2016). The marginal likelihood can also be used to weigh posterior samples from different hypotheses in order to average them (Kamary *et al.* 2014; Robert 2016), but we have not done that here.

### Simulation Procedure

We use simulation to evaluate our approach with respect to accuracy in allelic series inference and error in haplotype effect estimation. In particular, we focus on performance in the absence of additional phylogenetic information, and the utility of including that additional prior information, with varying levels of accuracy, as a coalescent tree. In the Supplement, we consider prior selection for the allelic series and concentration parameter.

We iteratively simulate single-locus QTL for a MPP with J=8 founder haplotypes, and at each locus we assume known but varied coalescent phylogeny of the haplotypes. We use a fixed experiment size of *N* = 400 individuals, balanced with respect to haplotypes, and with known homozygous diplotype states, **D**. Rather than vary experiment size, we instead vary QTL effect size, as measured by the proportion of total phenotype variance explained by the QTL, 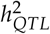. We anticipate that the results of these simulations depend primarily on power (a function of experiment size, haplotype balance, and QTL effect size), and thus hold both experiment size and haplotype balance fixed for simplicity. We also assume that the population does not have structure in genetic background, and without replicate observations, any variance attributable to strain effects is indistinguishable from individual-level error and can be ignored. We consider only additive QTL effects because the diplotypes are assumed to be homozygous in this simulation, and dominance effects would not be revealed.

Subject to these assumptions, the simulation procedure is as follows:

- Sample a coalescent tree *T* to describe the local phylogenetic relationship of the eight founder haplotypes:

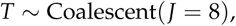
- For a given functional mutation rate *α*, calculate the distribution of allelic series implied by the tree:

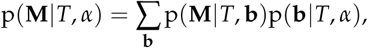
- For a given number of functional alleles *K*, sample an allelic series, conditional on *T*, that satisfies *K*:

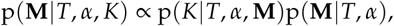
- For a given QTL effect size 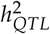, zero-center and scale *K* equally-spaced allele effects ***β***_alle_ to satisfy mean(***β***_alle_) = 0 and 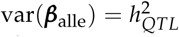
- Sample a vector of *N* individual errors from

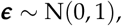

and zero-center and scale to satisfy mean(***ε***) = 0 and 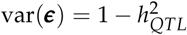,
- Compute the simulated phenotypes:

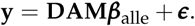

Rather than sample normally-distributed allele effects, we use uniformly-spaced effects, as in King *et al*. (2014). As the number of functional alleles increases, if the allele effects are normally-distributed, the minimum distance between any two allele effects becomes increasingly small, making it harder to distinguish these effects. Uniformly-spacing the effects eliminates the possibility of arbitrarily small and undetectable differences between alleles.

For each simulated experiment, we consider five possible scenarios, representing different levels of prior information about the allelic series and the underlying phylogeny:

- both the allelic series and coalescent tree are unknown (termed “CRP”),
- the allelic series is unknown and the coalescent tree is known (“Tree”),
- the allelic series is unknown and the coalescent tree is partially misspecified using the procedure defined in Azim Ansari and Didelot (2016) (“Misspecified”),
- the allelic series is unknown and the coalescent tree is completely misspecified as an unrelated coalescent tree (“Incorrect”), and
- an oracle approach where the allelic series is known (“Known”).

These are also compared with the standard haplotype-based approach, which assumes that all haplotypes are functionally distinct (“Full”).

The above scenarios are evaluated with respect to their accuracy in identifying the allelic series and their error in estimating haplotype effects. Specifically considered are:

- whether or not the MAP allelic series is the correct allelic series (“0-1 Accuracy”),
- the posterior mass on the correct allelic series (“Posterior Certainty”), and
- the mean squared error of the posterior haplotype effects relative to the true effects, averaged over posterior samples (“MSE”).

We perform 1000 simulations for each combination of the following parameter settings:

- Number of functional alleles *K*: [1-8],
- QTL effect size 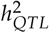 : [10%, 50%],
- Concentration parameter *α*: [1].

In all cases, results are stratified by the true number of functional alleles *K*. For this reason, our results are relatively insensitive to *α*, but we note that the relative benefit of tree information (conditional on *K*) is increased with higher *α* and decreased with lower *α*.

We do not directly compare our approach with other methods due to their considerable differences. Yalcin *et al*. (2005) consider only biallelic series and single variants. Mosedale *et al*. (2019) consider multiallelic series but only in the context of single variants. King *et al*. (2014) consider multiallelic series but only a subset of all multiallelic possibilities. Jannink and Wu (2003) consider all multiallelic series but do not include prior information about haplotype relatedness. Azim Ansari and Didelot (2016) include prior information about haplotype relatedness but consider categorical rather than quantitative traits. Given these differences, we discuss our findings in the context of these other methods rather than attempt a direct comparison.

### Data

We apply our allele-based association approach to three real-data examples, each of which highlights a key point about our approach for allelic series inference. The first example, an analysis of a QTL for a red blood cell phenotype detected in the PreCC by Kelada *et al*. (2012), introduces allelic series inference and demonstrates how it is improved by information on the local phylogeny. The second example, an analysis of whole lung cis-eQTL detected in the PreCC by Kelada *et al*. (2014), summarizes the distribution of allelic series over many QTL and identifies QTL that appear highly multiallelic. These separate studies used many of the same mice but analyzed different phenotypes. The third example, an analysis of two whole head cis-eQTL detected in the DSPR by King *et al*. (2014), shows that our allele-based approach (without tree information) is applicable even when there are many founder haplotypes (*J* = 15 instead of *J* = 8). These examples are detailed below.

#### Phenotypes

Our first example uses data from Kelada *et al*. (2012), a study of of blood parameters in *N* = 131 PreCC mice. This study identified a large-effect QTL for mean red blood cell volume (MCV) on chromosome 7 and a candidate causal gene, *Hbb-bs*, for the QTL.

Our second example uses data from Kelada *et al*. (2014), a study of whole-lung gene expression in *N* = 138 PreCC mice. Gene expression was measured by microarray and rank-normalized prior to eQTL mapping. For our analyses, we focused on 4,509 genes with cis-eQTL (within 10Mb of the gene), and we ignored eQTL for which array probes contained SNPs segregating between the founder strains, as these bias the microarray and are a potential source of false positive QTL (Alberts *et al.* 2007).

Our third example uses data from King *et al*. (2014), a study of whole-head gene expression in *N* = 596 crosses of DSPR fly lines. Gene expression was measured by microarray and rank-normalized prior to eQTL mapping. The authors highlighted two examples, CG4086 and CG10245, as examples of biallelic and multiallelic eQTL, respectively. We focus on these two examples for our analyses.

#### Diplotypes

The PreCC studies did not report the full diplotype state probabilities that are required for our approach, only additive haplotype dosages. We therefore performed another haplotype reconstruction using the published genotype information, also using HAPPY (Mott *et al.* 2000) and assuming a genotyping error rate of 0.01 as in Aylor *et al*. (2011). The studies averaged haplotype dosages from adjacent loci if there was no evidence of recombination across them in the PreCC population. To remain consistent with the published results, we averaged the diplotype state probabilities from our new haplotype reconstruction over the same regions.

The DSPR is comprised of two separate *J* = 8 populations with one shared founder, for a total of *J* = 15 founder haplotypes. The data we analyze are crosses of lines from the two separate populations. King *et al*. (2012) report full 36-state (homozygous and heterozygous) diplotype probabilities for all lines in the two separate *J* = 8 populations. To compute diplotype state probabilities for crosses of lines, we enumerated all possible combinations of diplotype states in the crossing lines, calculated the probability of possible haplotype combinations in the resulting cross, and weighed these combinations by the diplotype state probabilities of the lines. We assumed that maternal and paternal copies of the shared founder haplotype were identical. This results in full diplotype state probabilities for all crosses of lines and accounts for possible heterozygosity in both the lines and the crosses.

#### Phylogeny

For the MCV QTL analysis, we assumed that *Hbb-bs* is causal and inferred the phylogenetic tree of the founder haplotypes at this genomic region. First, we identified the location of *Hbb-bs* (Chr7: 103,826,523-103,827,928 in GRCm38/mm10), as reported by Mouse Genome Informatics (Bult *et al.* 2019). Next, we identified a larger 23kb nonrecombinant region surrounding the gene (Chr7: 103,807,679 103,831,178) by applying the four-gamete test (Hudson and Kaplan 1985) to high-quality SNPs from Sanger Mouse Genomes Project Keane *et al*. (2011). Then, we constructed a sequence alignment for the founder haplotypes using high quality SNPs and indels from the same source. Next, we used BEAST 1.8.3 (Drummond *et al.* 2012) to infer a coalescent phylogeny for this sequence alignment, assuming a constant mutation rate, constant population size and the HKY substitution model (Hasegawa *et al.* 1985). We generated one million Markov chain Monte Carlo (MCMC) samples from the posterior of coalescent trees, thinning every 1000 samples, yielding a total of 1000 posterior samples of the tree. These trees are visualized using Densitree (Bouckaert and Heled 2014). Lastly, we computed the allelic series prior distribution for each sample of the tree and averaged the results in order to arrive at a final tree-informed allelic series prior distribution for this QTL.

For both sets of eQTL analyses, we assumed that phylogenetic information for each QTL was unknown and used the CRP prior distribution for the allelic series, which implicitly integrates over all coalescent phylogenies.

### Computation

Posterior sampling proceeded by drawing 100,000 samples from a single MCMC chain for each analysis, with results reported based on the entire chain. The only exceptions were the DSPR CRP analyses, where we drew 1,000,000 samples due to the larger space of possible allelic series in the DSPR. We note that stable results were obtained for most analyses using 1/10 the number of samples, and we expect that fewer samples will be sufficient for many applications.

The MCV and DSPR analyses were run in Microsoft R Open 3.5.3 on an ASUS G10AJ-US010S desktop computer with Intel Core i7-4790 (3.6GHz) processor and 16GB of RAM. Computation time for these analyses is in **Table S1**. The the other analyses were run in parallel using a distributed computing cluster (https://its.unc.edu/research-computing/longleaf-cluster/), and their computation time is not reported due to varying hardware.

### Availability of Software and Data

These methods are implemented in the Tree-Based Inference of Multiallelism via Bayesian Regression ‘TIMBR’ R package, available on GitHub at https://github.com/wesleycrouse/TIMBR. The simulations and data are available in an accompanying repository at https://github.com/wesleycrouse/TIMBR_data.

## Results

### Simulation: Allelic Series Accuracy and Haplotype Effect Estimation

Based on our simulations, the accuracy of allelic series inference depends on three factors: 1) the true number of functional alleles, 2) the QTL effect size, and 3) the level of prior information about the phlyogenetic tree. Accuracy is defined here in the 0-1 sense, as whether the allelic series with the greatest posterior probability (the MAP) is correct. This is shown in **Figure 3** for different levels of tree information, numbers of true functional alleles, and effect sizes. In the high power scenario (**Figure 3B**), allelic series inference without tree information is accurate for as many as *K* = 5 alleles. In the low power scenario (**Figure 3A**), however, it is possible to accurately identify biallelic but not multiallelic series. Across both scenarios, additional tree information provides a modest increase in accuracy, even when that information is partially misspecified. Including completely misspecified tree information substantially decreases allelic series accuracy.

**Figure 3.**
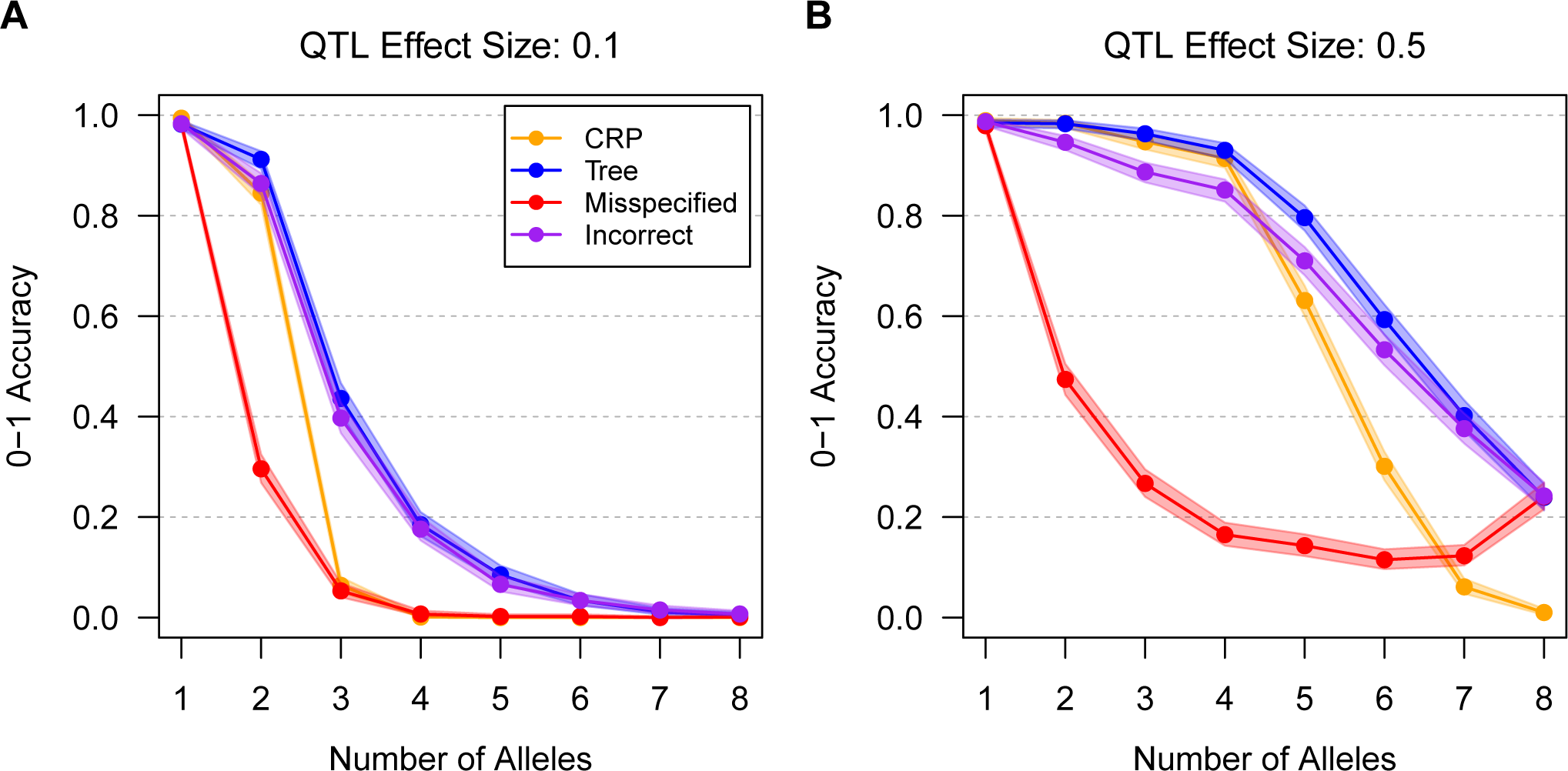
0-1 accuracy of posterior allelic series inference with four different levels of tree information, for varying numbers of true functional alleles, across two effect sizes. CRP assumes no tree information, Tree assumes perfect tree information, Misspecified assumes a partially misspecified tree, and Incorrect assumes a completely misspecified tree. Points are connected for clarity. Shading denotes 95% confidence intervals. In the low power scenario (**A**), accuracy for the CRP is high when the QTL has two or fewer alleles but low when it is multiallelic. In the high power scenario (**B**), accuracy for the CRP is high for an intermediate number of alleles but low when there are many alleles. Across both scenarios, Tree information provides a modest increase in accuracy relative to the CRP, even when that information is Misspecified. Incorrect tree information decreases allelic series accuracy.

Related to, but distinct from, accuracy is posterior certainty, or the posterior probability of the correct allelic series (**Figure 4**). This may be low even when accuracy is high. In the high power scenario (**Figure 4B**), allelic series inference without tree information is highly certain when there are *K* = 4 or fewer alleles, but the posterior mass is less than 50% when there are more alleles. In the low power scenario (**Figure 4A**), all multiallelic series are uncertain. Tree information increases certainty even when partially misspecified. Notably, posterior certainty when the tree is completely misspecified is similar to posterior certainty with no tree information, despite reducing 0-1 accuracy. This suggests that, on average, incorrect tree information increases certainty on incorrect allelic series, rather than reducing certainty on the correct allelic series (although the latter must also occur, when the incorrect tree does not permit the correct allelic series).

**Figure 4.**
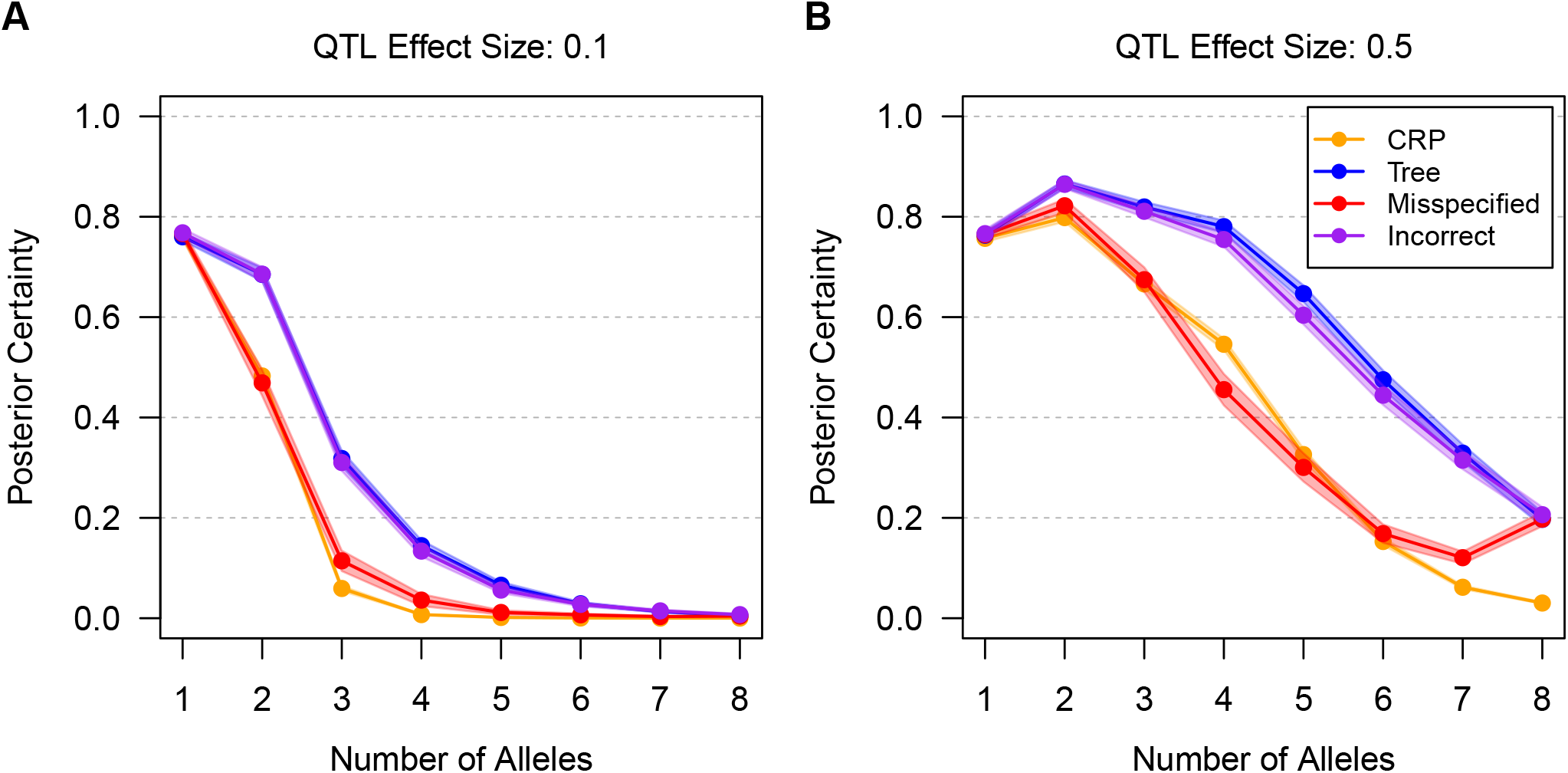
Posterior certainty of the correct allelic allelic series with four different levels of tree information, for varying numbers of true functional alleles, across two effect sizes. CRP assumes no tree information, Tree assumes perfect tree information, Misspecified assumes a partially misspecified tree, and Incorrect assumes a completely misspecified tree. Points are connected for clarity. Shading denotes 95% confidence intervals. In the low power scenario (**A**), posterior certainty for the CRP is high when the QTL has two or fewer alleles but low when it is multiallelic. In the high power scenario (**B**), posterior certainty is high for an intermediate number of alleles but low when there are many alleles. Across both scenarios, Tree information increases posterior certainty relative to the CRP, even when that information is Misspecified. Incorrect tree information has similar accuracy to the CRP on average.

In the Supplement, we show that there is a general tendency to underestimate the number of alleles when the true number of alleles is high. This is consistent with simulation results in both Azim Ansari and Didelot (2016) and King *et al*. (2014).

Despite uncertainty in the allelic series, the allele-based association approach can improve haplotype effect estimation. **Figure 5** shows the MSE of haplotype effect estimates. In the high power scenario (**Figure 5B**), the tree-naive allele-based approach has lower MSE than the haplotype-based approach, provided there are fewer than *K* = 5 functional alleles at the QTL. Including prior tree information improves MSE relative to the tree-naive case, even when the tree is partially misspecified. Including completely misspecified tree information increases MSE relative to the tree-naive case, although it still outperforms the haplotype-based approach when there are an intermediate number of alleles. Results for the low power scenario (**Figure 5A**) are similar, though the allele-based approaches are generally less beneficial when QTL are multiallelic.

**Figure 5.**
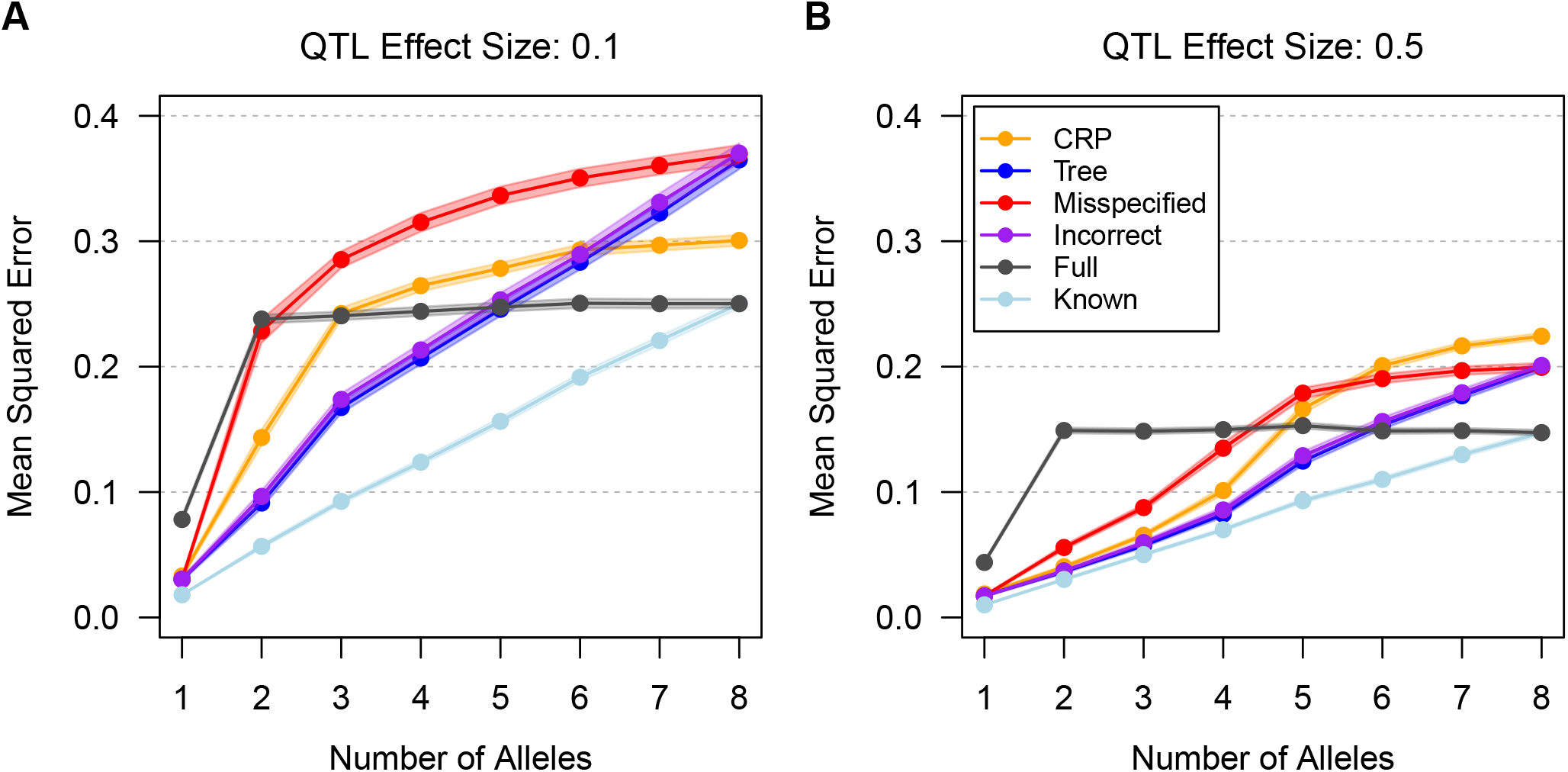
Mean squared error (MSE) of haplotype effect estimates with four different levels of tree information, for varying numbers of true functional alleles, across two effect sizes. CRP assumes no tree information, Tree assumes perfect tree information, Misspecified assumes a partially misspecified tree, and Incorrect assumes a completely misspecified tree. Full is the haplotype-based approach where all haplotypes are functionally distinct, and Known is an oracle prior in which the correct allelic series is known. Points are connected for clarity. Shading denotes 95% confidence intervals. In the low power scenario (**A**), CRP has lower MSE than Full when the QTL has two or fewer alleles. In the high power scenario (**B**), CRP has lower MSE than Full for an inter-mediate number of alleles. Across both scenarios, Tree information improves MSE relative to CRP, even when it is Misspecified, and it outperforms Full for an intermediate number of alleles. Incorrect tree information increases MSE relative to CRP, but it still outperforms Full for an intermediate number of alleles.

To summarize, these results suggest that, in practice, there will often be considerable uncertainty in allelic series inference, especially when effect sizes are small and QTL are multiallelic. Nonetheless, accounting for the allelic series can still improve haplotype effect estimation relative to the standard haplotype-based approach. These finding are consistent with simulation results in Jannink and Wu (2003). In the Supplement, we present additional simulations that guide prior specification and provide further insight into allelic series inference.

### Example One: Allelic Series Inference with Tree Information

We analyzed the MCV QTL previously identified in the PreCC study of Kelada *et al*. (2012). **Figure 6A** shows the MCV phenotype for the 94 of 131 mice with prior maximum diplotype states that are homozygous at the QTL, plotted by that haplotype. Heterozygous mice are omitted to simplify the figure, but they are included (with diplotype uncertainty for all mice) in subsequent analyses. Based on the figure, the phenotype clearly depends on the haplotype at the QTL, but intuitively, the number of functional alleles is not obvious.

**Figure 6.**
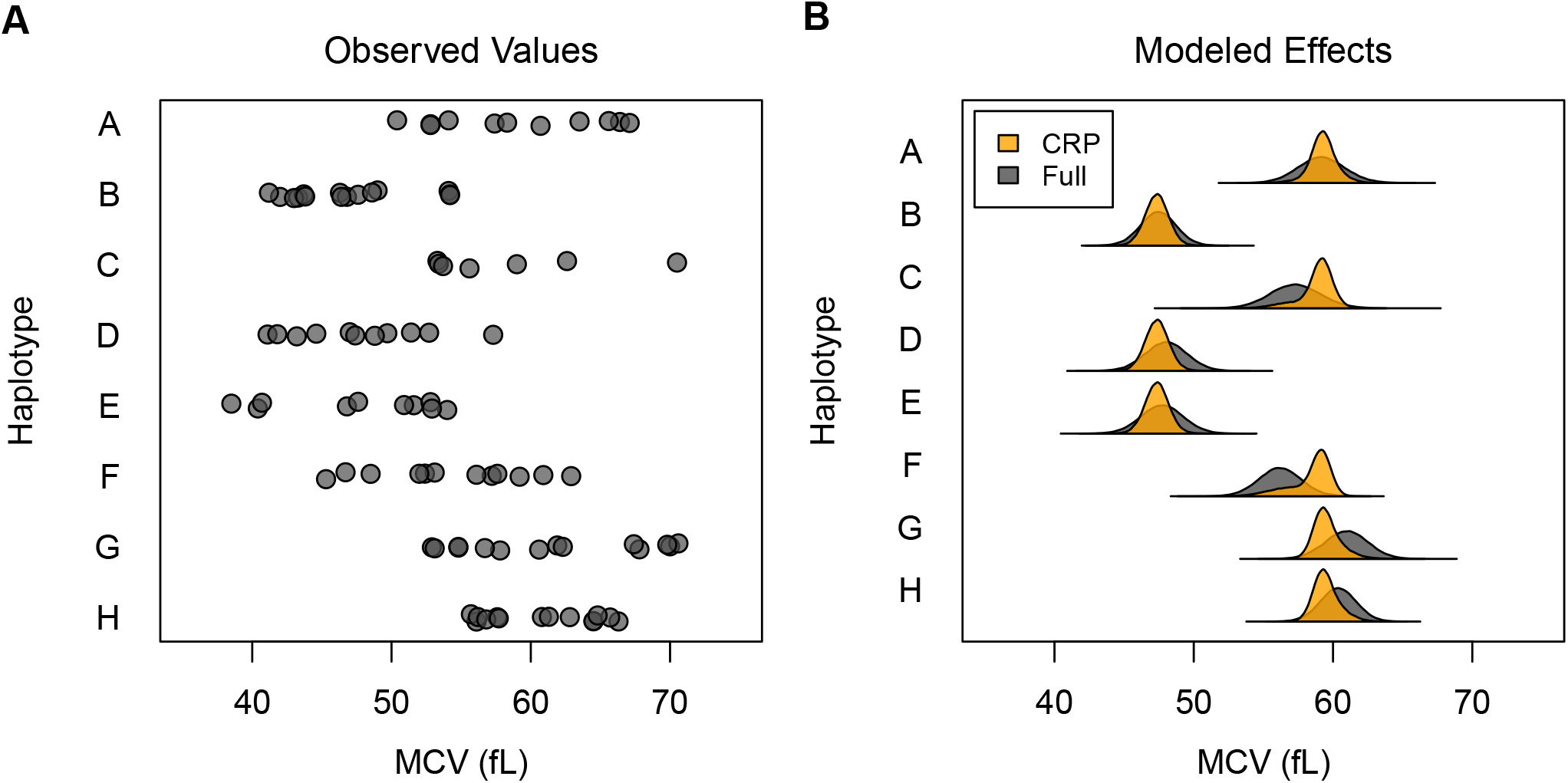
Mean Cell Volume (MCV) in the PreCC by founder haplotype at the QTL, and posterior distribution of haplotype effects using the Full and CRP approaches. (**A**) MCV by founder haplotype for mice with homozygous prior maximum diplotype state. Prior maximum diplotype state is homozygous for 94 of 131 mice. Heterozygous mice are omitted for clarity, but they are included (with diplotype uncertainty for all mice) in other analyses. The phenotype depends on the haplotype at the QTL, but intuitively, the number of functional alleles is not obvious. (**B**) Posterior distribution of haplotype effects using the Full and CRP approaches. Full is the haplotype-based approach where all haplotypes are functionally distinct, and CRP assumes no tree information. CRP haplotype effects are more certain than Full because the allelic series allows information about effects to be shared across haplotypes. This is evident for haplotype F, which has its effect distribution pulled towards an allele effect that is shared with haplotypes A, C, G, and H.

The top ten posterior allelic series inferred using the CRP approach are shown in **Table 1**. The top allelic series is biallelic and comprises 55.7% of the posterior probability, but there are several other multiallelic series with reasonable support (11 with ≥1%, together accounting for another 30.1% of the posterior). These multiallelic series preserve the biallelic contrast identified by the top allelic series, indicating that this is a high-confidence feature of the haplotype effects. The posterior distribution of the number of alleles is given in **Figure 7**, and the posterior expected number of alleles is 2.59. Overall, these results provide evidence in favor of a biallelic QTL, but allelic series inference is still uncertain.

**Table 1.** Top ten posterior allelic series for the MCV QTL in the PreCC using the CRP approach.

|  | Allelic Series | # of Alleles | Posterior Probability |
| --- | --- | --- | --- |
| 1 | 0,1,0,1,1,0,0,0 | 2 | 0.5568 |
| 2 | 0,1,0,1,1,2,0,0 | 3 | 0.0801 |
| 3 | 0,1,2,1,1,2,0,0 | 3 | 0.0644 |
| 4 | 0,1,0,1,1,0,2,2 | 3 | 0.0278 |
| 5 | 0,1,0,1,1,0,2,0 | 3 | 0.0204 |
| 6 | 0,1,0,2,1,0,0,0 | 3 | 0.0189 |
| 7 | 0,1,2,1,1,0,0,0 | 3 | 0.0181 |
| 8 | 0,1,0,1,2,0,0,0 | 3 | 0.0178 |
| 9 | 0,1,0,2,2,0,0,0 | 3 | 0.0160 |
| 10 | 0,1,0,1,1,0,0,2 | 3 | 0.0132 |

**Figure 7.**
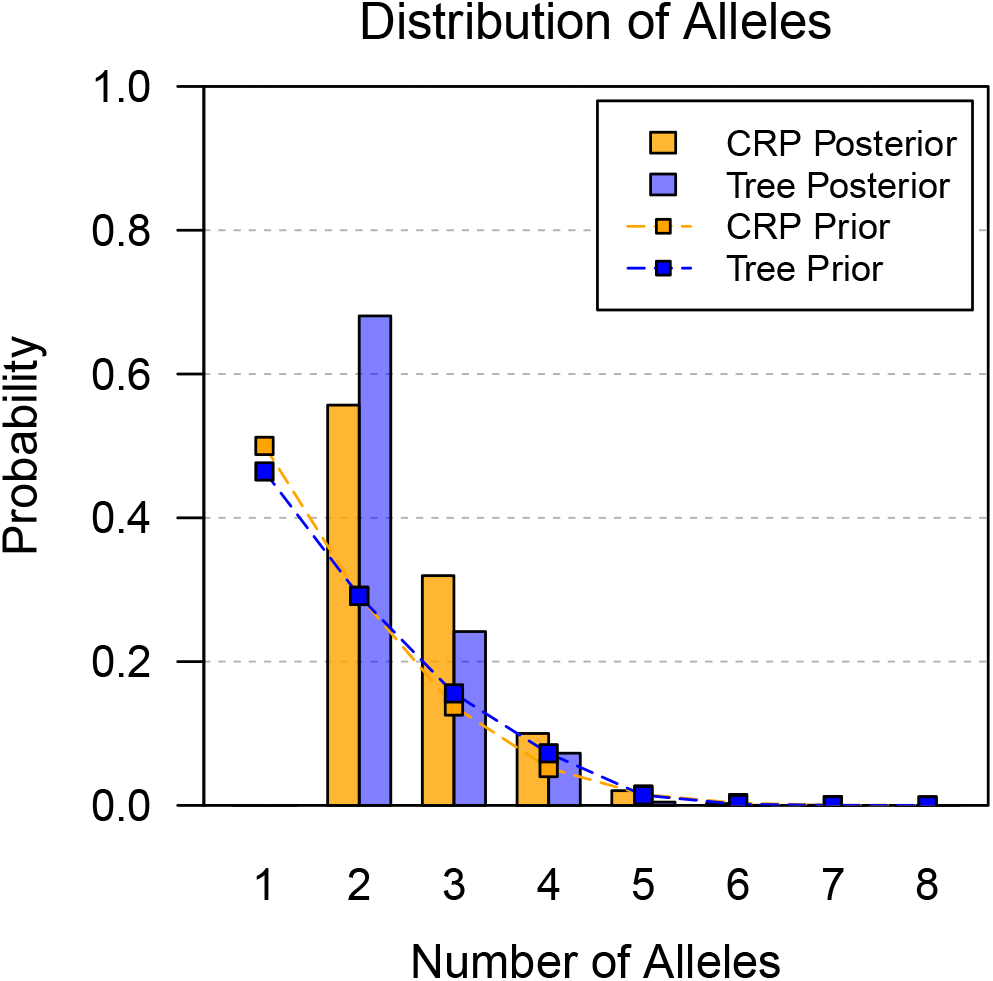
Posterior distribution of number of alleles for the MCV QTL in the PreCC using the CRP and Tree approaches. CRP assumes no tree information, and Tree assumes tree information averaged over 1000 tree samples. Connected points denote the corresponding prior distributions. The prior distributions favor smaller numbers of alleles. The posterior probability of a biallelic QTL is high using CRP, and it is further increased by Tree.

The posterior distribution of haplotype effects using both the Full and CRP approaches is shown in **Figure 6B**. As expected, the Full haplotype effect estimates are similar to the observed phenotypes. Relative to the Full, the CRP haplotype effects are more certain, with narrower 95% highest posterior density (HPD) intervals, as shown in **Table S2**. This increased certainty is because the allelic series model allows information about the effects to be shared across haplotypes. This is particularly evident for haplotype F, which has its effect distribution pulled towards an allele effect that is shared with haplotypes A, C, G, and H. Nonetheless, the haplotype effect distribution of F has a long tail, covering much of the original range of the Full haplotype effect distribution. Comparing the Full and CRP approaches more broadly, the lnBF in favor of the CRP is 1.17, indicating positive evidence in favor of allele-based effects.

Samples of phylogenetic trees that relate the founder haplotypes at the causal locus are shown in **Figure 8B**. In general, there are long branches separating haplotypes B, D, and E from the other five haplotypes. Among the remaining haplotypes, A, H, and C are also more closely related than F and G. Relative to the coalescent, (**Figure 8A**), these trees are highly structured, representing only 3 of 10,395 possible tree topologies. This informs the prior distribution of the allelic series in the Tree model (**Figure 8C-D** and **Table S3**). There are 720 allelic series with support using the Tree approach, compared with the full space of 4,140 using the CRP. The allelic series favored by the Tree approach reflect the relationships encoded by the causal trees; for example, the top non-null allelic series is biallelic and contrasts haplotypes B, D, and E against the others, and its prior probability is increased over 150-fold relative to the CRP.

**Figure 8.**
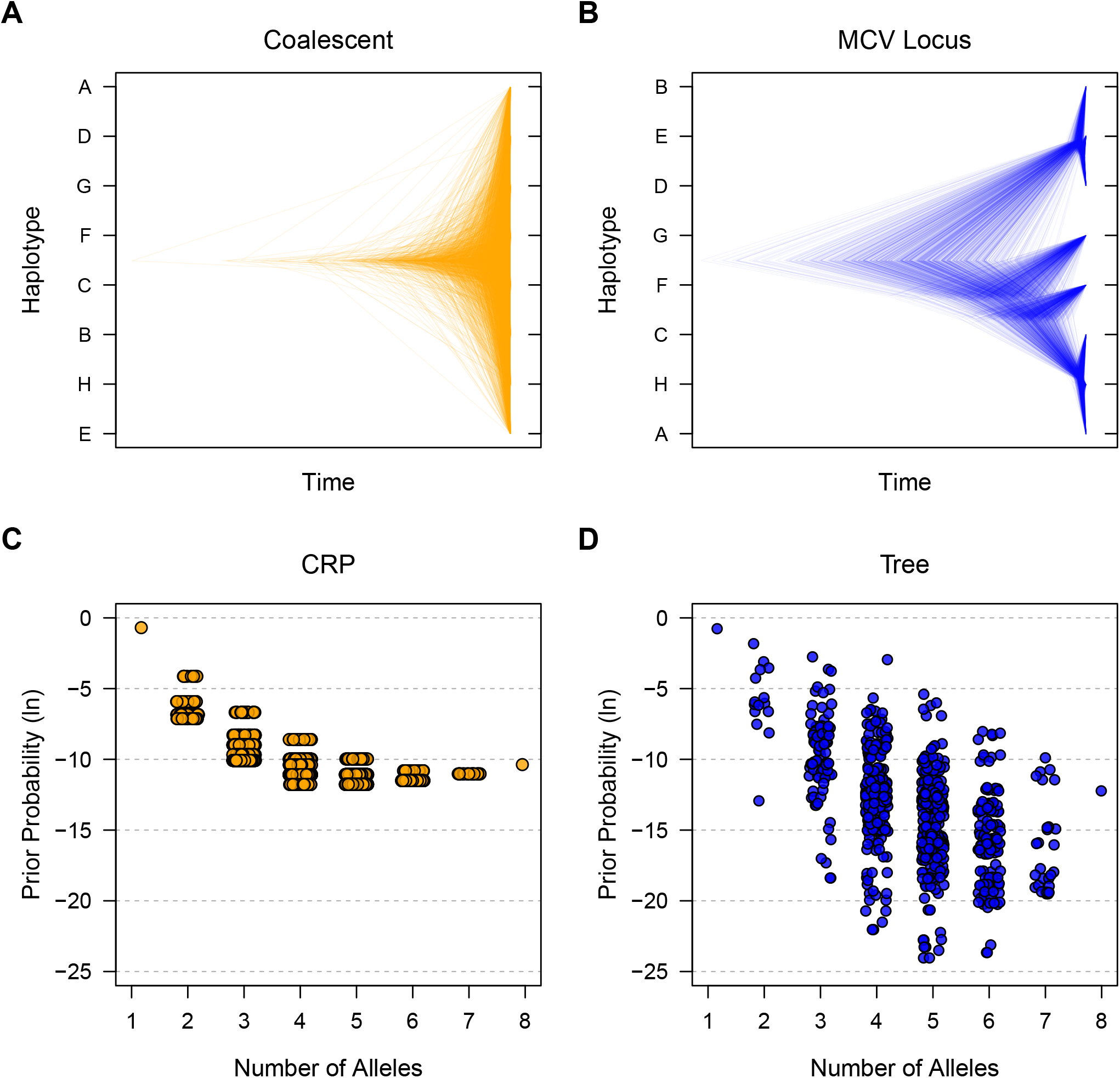
Prior distributions of trees and allelic series for the MCV QTL in the PreCC using the CRP and Tree approaches. (**A**) 1000 samples of random coalescent trees. This is the implied distribution over trees for CRP. The trees are unstructured, with 10,395 different topologies that could be sampled. (**B**) 1000 samples of trees at the MCV causal locus. This is the prior distribution over trees for the Tree approach. The trees are highly structured, with only 3 different topologies sampled. Long branches separate haplotypes B, D, and E from the other haplotypes. (**C**) Prior distribution of allelic series by number of alleles using CRP. There are 4,140 possible allelic series, with many having identical prior probability. (**D**) Prior distribution of allelic series by number of alleles using CRP. There are 720 possible allelic series, with prior probability reflecting the relationships encoded by the causal trees. Tree information informs the allelic series prior distribution relative to CRP.

The top ten posterior allelic series inferred using the Tree approach are shown in **Table 2**. The top allelic series is unchanged from the CRP results, but its posterior probability is increased to 68.1%. There are fewer multiallelic series with reasonable support (5 with ≥1% posterior probability), and they have been informed by the phylogenetic distance between F, G and the other haplotypes. The posterior distribution of the number of alleles is given in **Figure 7**, and the posterior expected number of alleles is 2.40.

**Table 2.** Top ten posterior allelic series for MCV QTL in the PreCC using the Tree approach.

|  | Allelic Series | # of Alleles | Posterior Probability |
| --- | --- | --- | --- |
| 1 | 0,1,0,1,1,0,0,0 | 2 | 0.6808 |
| 2 | 0,1,0,1,1,2,0,0 | 3 | 0.1335 |
| 3 | 0,1,0,1,1,0,2,0 | 3 | 0.0865 |
| 4 | 0,1,0,1,1,2,3,0 | 4 | 0.0639 |
| 5 | 0,1,0,1,1,2,2,0 | 3 | 0.0122 |
| 6 | 0,1,2,1,1,0,0,0 | 3 | 0.0027 |
| 7 | 0,1,0,2,2,0,0,0 | 3 | 0.0027 |
| 8 | 0,1,2,1,1,3,0,0 | 4 | 0.0021 |
| 9 | 0,1,2,1,1,3,4,0 | 5 | 0.0019 |
| 10 | 0,1,0,1,1,0,0,2 | 3 | 0.0015 |

The posterior distribution of Tree-informed haplotype effects are largely unchanged from the CRP haplotype effects and are not shown for this reason. Overall, there is strong positive evidence for the Tree approach relative to the CRP, with a lnBF of 4.81 in favor of the Tree.

In summary, this example demonstrates that our method can be used to infer the allelic series at a QTL, that it can improve haplotype effect estimation, and that including additional phylogenetic information can increase the posterior certainty of the allelic series.

### Example Two: Identifying Multiallelic QTL

We analyzed the lung cis-eQTL previously identified in the PreCC study of Kelada *et al*. (2014). **Figure 9** shows the posterior distribution of number of alleles, averaged over all cis-eQTL. This suggests that many eQTL are multiallelic, with 35.7% and 21.8% posterior probability for *K* = 3 and *K* = 4 alleles, respectively. Given that the CC founders are comprised of three different subspecies of mice (Didion and Pardo-Manuel de Villena 2013), this multiallelism is reasonable. There is also substantial support for *K* = 2 biallelic eQTL, which has 30.3% posterior probability. These QTL were genome-wide significant when detected, so it is not surprising that there is near-zero support for the null model of *K* = 1 allele. There are also genes that appear highly multiallelic. **Table 3** highlights the most highly multiallelic QTL in this dataset.

**Table 3.** Highly multiallelic cis-eQTL for lung expression in the PreCC; top twenty by posterior expected number of alleles. Gene positions from NCBI37/mm9.

|  | Probe | Gene | Chr | Position | Expected Alleles |
| --- | --- | --- | --- | --- | --- |
| 1 | ILMN_2667352 | Glo1 | 17 | 30,729,806 | 6.7549 |
| 2 | ILMN_2418957 | Tmem181b-ps | 17 | 6,270,475 | 6.3900 |
| 3 | ILMN_3023451 | Zfp985 | 4 | 146,918,112 | 6.3694 |
| 4 | ILMN_2880052 | Xlr4b | X | 70,459,704 | 6.2664 |
| 5 | ILMN_2499598 | 2310058D17Rik | 11 | 58,777,283 | 6.1449 |
| 6 | ILMN_3004949 | Fam55d | 9 | 47,970,198 | 5.8813 |
| 7 | ILMN_2643495 | Fez1 | 9 | 36,640,394 | 5.8658 |
| 8 | ILMN_2685581 | H2-K1 | 17 | 34,132,957 | 5.7066 |
| 9 | ILMN_1221376 | Cyp4f39 | 17 | 32,589,668 | 5.6539 |
| 10 | ILMN_3153940 | Unc45b | 11 | 82,724,831 | 5.6496 |
| 11 | ILMN_2634905 | Fbp2 | 13 | 62,938,245 | 5.6022 |
| 12 | ILMN_2998406 | Zfp979 | 4 | 146,986,048 | 5.5955 |
| 13 | ILMN_1213056 | Fez1 | 9 | 36,640,394 | 5.5861 |
| 14 | ILMN_1236008 | Isoc2a | 7 | 4,828,740 | 5.5309 |
| 15 | ILMN_2776728 | Zfp979 | 4 | 146,986,048 | 5.4501 |
| 16 | ILMN_2735046 | Cml3 | 6 | 85,711,089 | 5.4235 |
| 17 | ILMN_2527805 | Wfdc10 | 2 | 164,481,546 | 5.4121 |
| 18 | ILMN_2665266 | H2-T22 | 17 | 36,175,354 | 5.4084 |
| 19 | ILMN_2894678 | H2-T22 | 17 | 36,175,354 | 5.4022 |
| 20 | ILMN_2584887 | Atp5f1 | 3 | 105,745,781 | 5.3849 |

**Figure 9.**
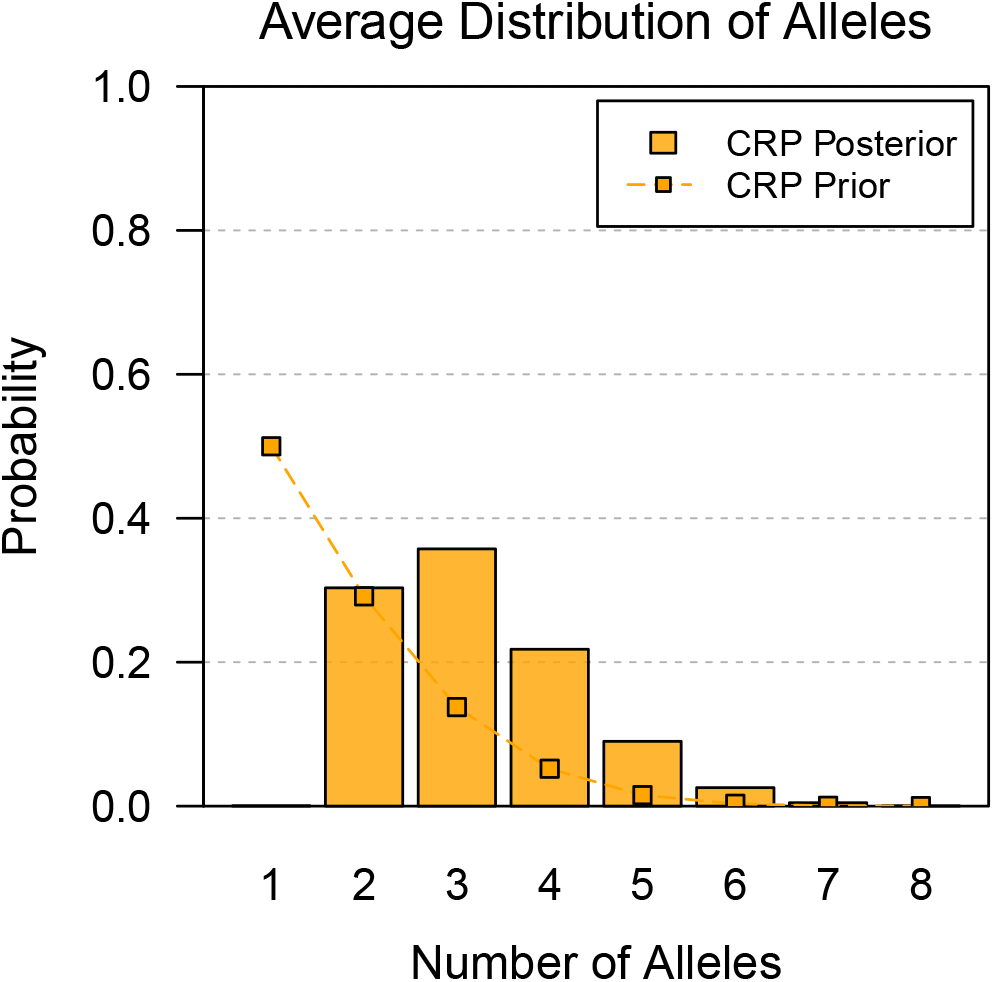
Posterior distribution of number of alleles, averaged over all cis-eQTL identified in whole lung expression in the PreCC. Connected points denote the corresponding prior distribution. The prior distribution favors smaller numbers of alleles. The posterior distribution is concentrated between two and four alleles, with considerable support for multiallelic series.

The most multiallelic cis-eQTL in our dataset was *Glo1*. **Figure 10A** shows *Glo1* expression for the 111 of 138 mice with prior maximum diplotype states that are homozygous at the QTL, plotted by that haplotype. As before, heterozygous mice are omitted to simplify the figure, but they are included during analysis. Our approach finds over 95% posterior support for six to eight alleles at this QTL (**Figure 10B**). Interestingly, previous studies have found that mouse strains have a complicated haplotype structure with many functional alleles at *Glo1*, and that expression of this gene is associated with anxiety-like behavior in mice (Williams IV *et al.* 2009). This supports our finding that *Glo1* is highly multiallelic.

**Figure 10.**
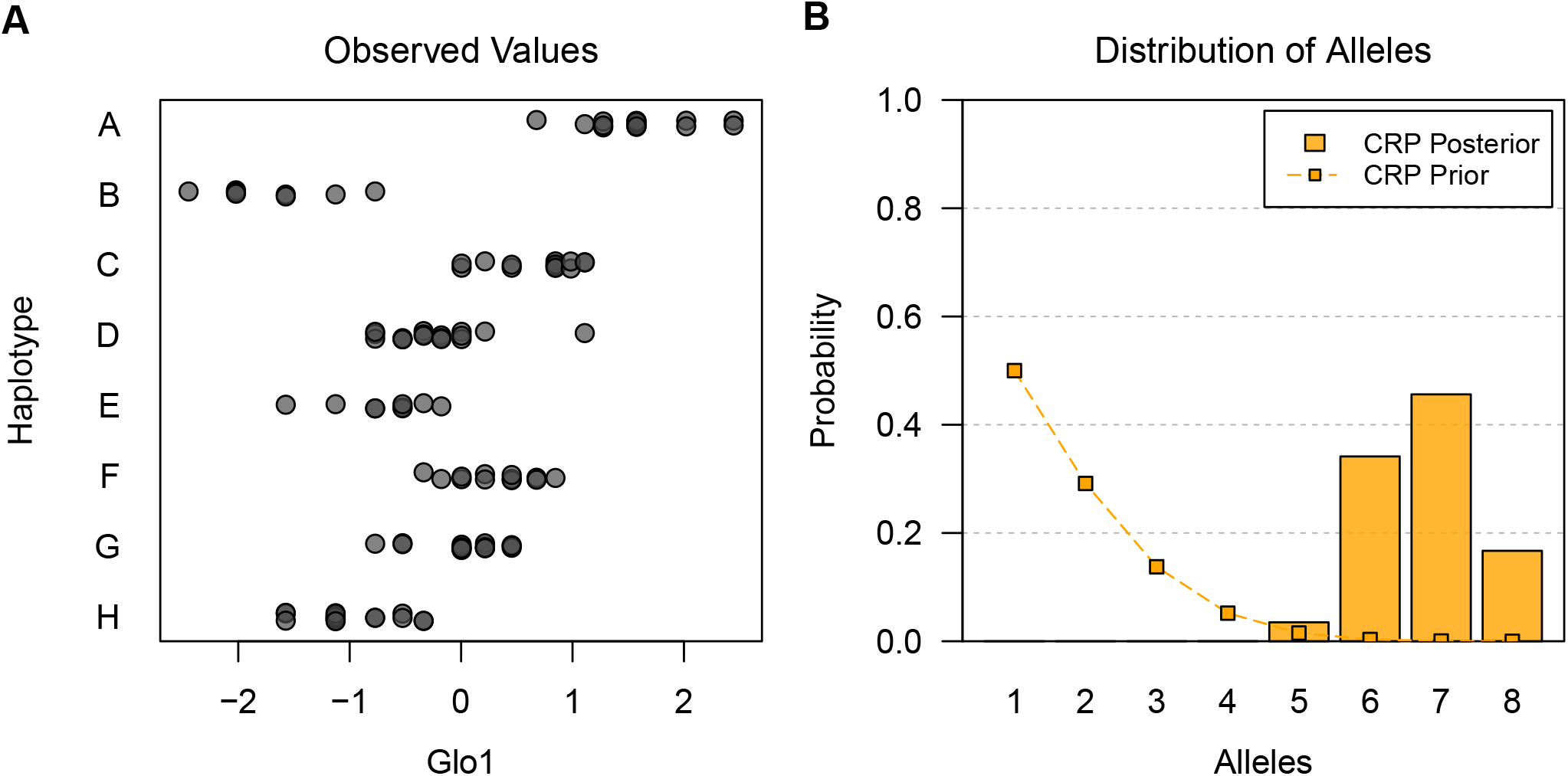
*Glo1* expression in the PreCC by founder haplotype at the QTL and posterior distribution of number of alleles. (**A**) *Glo1* expression by founder haplotype for mice with homozygous prior maximum diplotype state. Prior maximum diplotype state is homozygous for 111 of 138 mice. Heterozygous mice are omitted for clarity, but they are included (with diplotype uncertainty for all mice) in other analyses. The QTL appears highly multiallelic. (**B**) Posterior distribution of number of alleles for *Glo1* cis-eQTL. Connected points denote the prior distribution. The prior distribution favors smaller numbers of alleles. The posterior distribution is concentrated between six and eight alleles, indicating that the QTL is highly multiallelic.

We also note that several highly multiallelic cis-eQTL are near the major histocompatibility complex on chromosome 17, which is consistent with high genetic diversity in this region (Lilue *et al*. 2019).

### Example Three: Allelic Series Inference with Many Founder Haplotypes

We analyzed two whole-head eQTL previously identified in the DSPR study of King *et al*. (2014). **Figure 11A** shows the posterior distribution of number of alleles for the CG4086 eQTL using the CRP approach. Although the previous study found that this eQTL was biallelic, we find a 61.7% posterior probability that the QTL has three functional alleles. **Table S4** shows the top ten posterior allelic series, which tend to contrast haplotypes A6, A7, and B2 against the others. This is consistent with the Full posterior haplotype effects, shown in **Figure 11B-C**. Relative to the Full, the allele-based haplotype effects of the CRP are more certain, with narrower 95% HPD intervals (**Table S5**). Notably, both the Full and CRP approaches make haplotype effect predictions for A1, A5, and B3, all of which are poorly represented at this QTL and were omitted in the previous study. Overall, there is very strong evidence in favor of the of the CRP relative to the Full approach, with a lnBF of 7.71.

**Figure 11.**
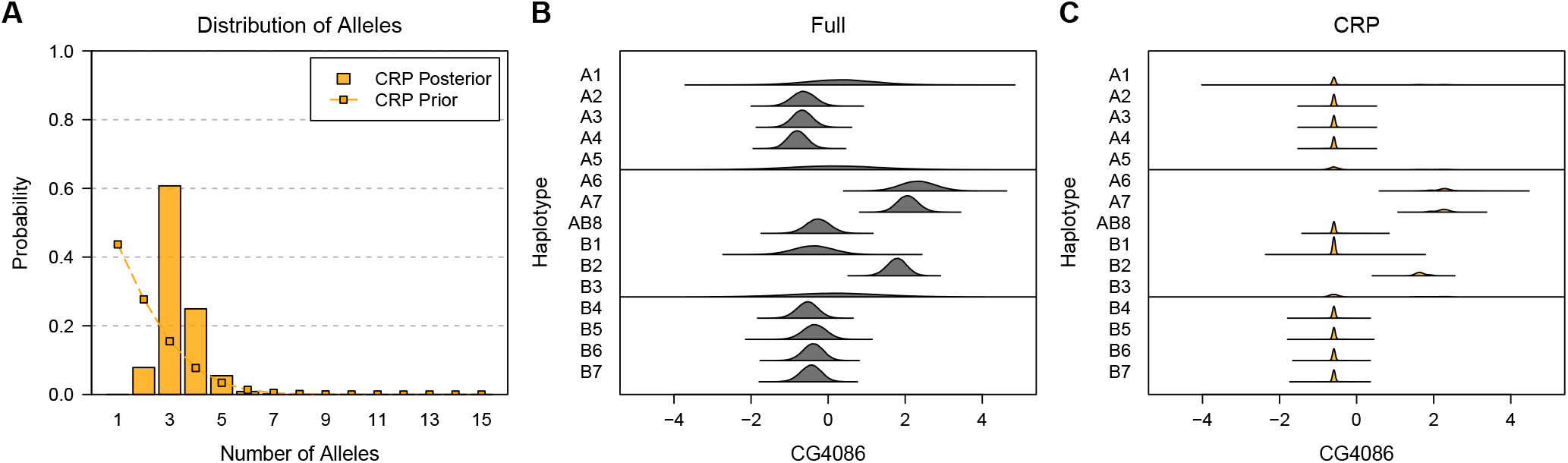
Posterior distribution of number of alleles and haplotype effects for the CG4086 cis-eQTL in the DSPR. (**A**) Posterior distribution of number of alleles. Connected points denote the prior distribution, which favors smaller numbers of alleles. The posterior distribution has considerable support for three alleles at this QTL. (**B**) Posterior distribution of haplotype effects using the Full approach. Full is the haplotype-based approach where all haplotypes are functionally distinct. (**C**) Posterior distribution of haplotype effects using the CRP approach. CRP assumes no tree information. CRP haplotype effects are more certain than Full because the allelic series allows information about effects to be shared across haplotypes. Both Full and CRP make effect predictions for A1, A5, and B3, all of which are poorly represented at this QTL.

The posterior distribution of number of alleles for the CG10245 eQTL using the CRP approach is shown in **Figure 12A**. The previous study found that this eQTL was highly multiallelic, a finding that we confirm, with an expected posterior number of alleles of 8.95. The posterior distribution of the allelic series, however, is highly uncertain (**Table S6**), due to the large number of possible allelic series when there are *J* = 15 founder haplotypes and many alleles. **Figure 12B-C** shows the posterior distribution of haplotype effects using the Full and CRP approaches. Interestingly, many of the haplotype effect distributions for the CRP are multimodal, and the 95% HPD intervals for the CRP are generally wider than for the Full (**Table S7**). This is a consequence of the highly uncertain posterior allelic series. The intervals for A4 and B2, both of which are poorly represented at this QTL, are actually narrower, showing how the CRP can provide, in a sense, additional shrinkage to the haplotype effects. Consistent with extensive multiallelism, there is very strong evidence against the CRP relative to the Full approach, with a lnBF of −11.15.

**Figure 12.**
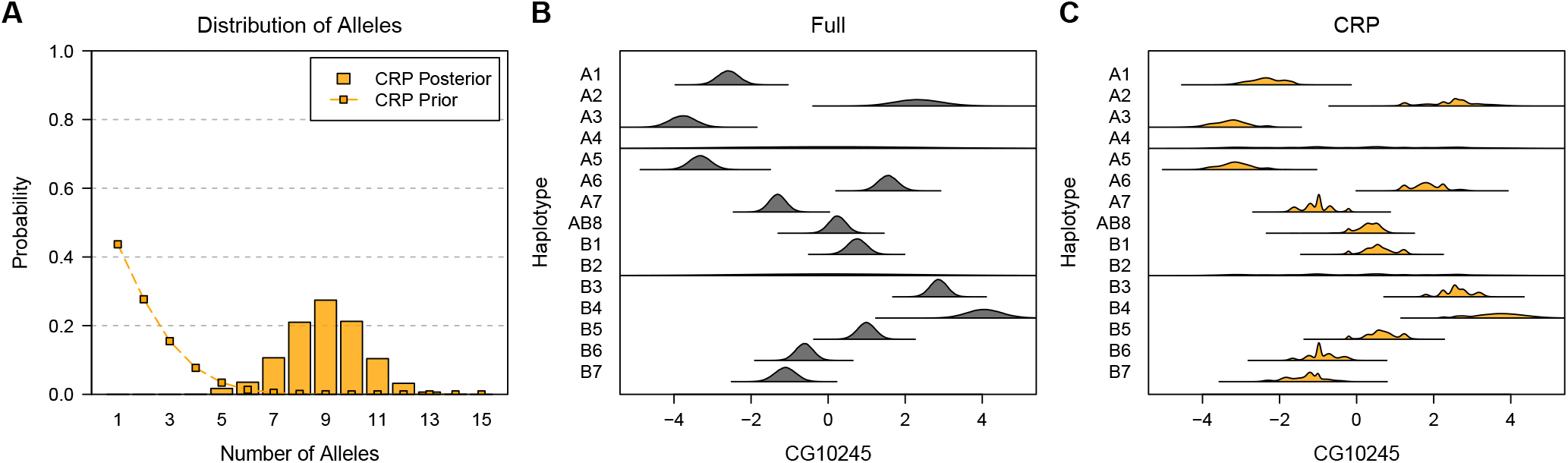
Posterior distribution of number of alleles and haplotype effects for the CG10245 cis-eQTL in the DSPR. (**A**) Posterior distribution of number of alleles. Connected points denote the prior distribution, which favors smaller numbers of alleles. The posterior distribution indicates the QTL is highly multiallelic. (**B**) Posterior distribution of haplotype effects using the Full approach. Full is the haplotype-based approach where all haplotypes are functionally distinct. (**C**) Posterior distribution of haplotype effects using the CRP approach. CRP assumes no tree information. CRP haplotype effects are multimodal due to the highly uncertain posterior allelic series. Both Full and CRP make effect predictions for A4 and B2, both of which are poorly represented at this QTL.

## Discussion

In this study, we developed a fully-Bayesian framework for inferring the allelic series in MPPs. Our approach partitions haplotypes into a potentially smaller number of functionally distinct alleles, and it accommodates prior information about haplotype relatedness. The allelic series is useful for investigating the genetic architecture of a QTL, and in particular, for determining if there are multiple causal variants at a locus. In this section, we summarize the findings of our study, suggest interesting directions for future research, and discuss the limitations of our approach.

### Allelic Series Inference is Uncertain but Useful

Our simulations indicate that inference of the allelic series (in the absence of tree information) is often uncertain, even in situations that we expect would have high QTL mapping power. This is especially true for multiallelic series. Posterior certainty is higher in the biallelic case, when there are relatively more observations to distinguish the allele effects. In combination with a prior distribution that expects few functional alleles, the posterior for biallelic series is decisive. Posterior certainty decreases, however, as the true number of functional alleles increases. This is because the space of possible allelic series configurations is larger (for an intermediate number of alleles), and there are relatively fewer observations to distinguish the allele effects. For these reasons, posterior certainty for multiallelic series is often quite low, unless effect sizes are very large or additional prior information can be reliably included.

In the Supplement, we considered a prior distribution that places more weight on higher numbers of alleles. This prior distribution had relatively better accuracy when the true number of alleles is high, but at the expense of decisive posterior certainty when the true number of alleles is low. For this reason, we focused our analyses on a parsimonious prior distribution that favors small numbers of functional alleles, though there may be scenarios in which a more permissive prior distribution is justified or desirable.

Taken together, these results suggest that, if multiallelism is common in MPPs, researchers will frequently find themselves following-up QTL with highly uncertain allelic series. In these cases, our allele-based association approach may be more useful for evaluating whether a QTL is more likely to be biallelic or multiallelic, rather than for identifying the allelic series *per se*. When the allelic series is uncertain, we recommend focusing on high-confidence features of the data rather than one specific allelic configuration. For example, although the posterior allelic series for the multiallelic QTL in the DSPR is extremely uncertain (**Figure 12C** and **Table S6**), it is still highly probable that haplotypes A1 and A3 are different than A2, even though they may not be the same. These inferences are straightforward in the allele-based framework, and they are often well-informed even when the allelic series is uncertain. Our approach can also be used comparatively, to determine the most multiallelic QTL in a dataset, as we did for lung cis-eQTL identified by Kelada *et al*. (2014). Characterizing highly multiallelic QTL is an interesting topic for future investigation, and this comparative inference does not require high certainty in the allelic series posterior.

### Allelic Series Inference Can Improve Effect Estimation

Despite uncertainty in the allelic series, the allele-based approach improves haplotype effect estimation relative to the haplotype-based approach, provided the locus has only a few functional alleles. The allele-based approach allows the data to be represented using fewer parameters, and this reduction in parameters can still be beneficial even when the allelic series is only partially known. This improvement in effect estimation was particularly evident in the “biallelic” DSPR example (**Table S5**). These results suggest that the allele-based approach will be useful in the context of phenotype prediction, or other applications that might benefit from improved effect estimation.

When there are many functional alleles at a locus, though, the allele-based approach increases error in haplotype effect estimation relative to the full haplotype-based approach. This is because our prior distribution favors small numbers of alleles, and, in the Supplement, we show that our approach tends to underestimate the number of alleles for highly multiallelic QTL. This biases haplotype effect estimates towards each other, and it can increase credible intervals for these effects (**Table S7**). In practice, it will useful to compare the fit of the allele-based and haplotype-based approaches using their BF. If there is decisive evidence in favor of the haplotype-based approach, it may be better to use the haplotype-based effect estimates. Another option is to weigh the posteriors of the haplotype-based and allele-based approaches using their marginal likelihoods, essentially placing a prior distribution over them (Kamary *et al.* 2014; Robert 2016). This avoids making a decision about which approach to use while still favoring the one which better describes the data.

We note that our simulations assumed uniformly-spaced allele effects. This assumption eliminates the possibility of arbitrarily small and undetectable differences between alleles, ensuring that alleles are “practically” distinct from one another. In real data, however, small differences between alleles may exist. If alleles effects were not uniformly spaced, our simulation results for *K* = 1 and *K* = 2 would be unchanged. For higher numbers of alleles, we expect that allelic series accuracy would be reduced but that MSE would be improved, as the model focuses on a smaller number of alleles that explain relatively more phenotypic variance. Characterizing our allele-based association approach under different genetic architectures is an interesting direction for further research.

### Local Phylogeny Improves Allelic Series Inference, Although is Itself Uncertain

Accounting for founder haplotype relatedness in an allele-based association framework is the primary innovation of our research. Our simulations show that including prior information about haplotype relatedness, in the form of a coalescent tree, improves our allele-based association approach with respect to allelic series inference and haplotype effect estimation.

The local phylogenetic tree of the founder haplotypes is necessarily unknown, however, and can only be observed indirectly through genetic variation. Our framework is based on the coalescent (Kingman 1982), which describes the phylogenetic relationship for a single nonrecombinant genomic region. The assumption of no recombination is necessary because, in recombinant systems, phylogeny varies throughout the genome due to incomplete lineage sorting (Degnan and Rosenberg 2009) and, particularly for the CC and DO founder strains, introgression (Yang *et al.* 2011; Didion and Pardo-Manuel de Villena 2013). This means that neighboring genomic regions can have distinct (but correlated) phylogenetic trees. The complex structure describing recombination events and varied local phylogeny is the “ancestral recombination graph” (ARG), and inferring it is the subject of active research (Rasmussen *et al.* 2014; Kelleher *et al*. 2019). If the ARG were known exactly, variation in haplotype phylogeny throughout the genome could be a useful source of information for allelic series inference and, perhaps, QTL mapping. In practice, though, the ARG will be uncertain, with regions that are poorly informed by mutations or biased due to errors. Due to this uncertainty, the inferred ARG will be less useful for QTL mapping than known local phylogeny, although it is unclear to what extent.

Given uncertainty in the ARG, we recommend using our CRP (*i*.*e*., tree-naive) approach by default when analyzing QTL in MPPs. There are situations, though, when local phylogeny can be accurately inferred, and in these cases, including tree information improves allelic series inference and haplotype effect estimation. We demonstrated this for the MCV QTL, which has a known causal gene (Kelada *et al.* 2012). We anticipate our tree-informed approach will be useful in haploid systems, as in Azim Ansari and Didelot (2016) and Cybis *et al*. (2018), because most haploids do not recombine, and thus have a single phylogenetic history for their entire genome.

### Connecting the Allelic Series to Causal Variants

The allele-based association approach is useful for evaluating whether a QTL is more likely to be biallelic or multiallelic. It can be difficult, however, to connect information about the allelic series to causal variants. Evaluating evidence in favor of a single biallelic or multiallelic variant is straightforward, as our framework encompasses a fully-Bayesian implementation of merge analysis (Yalcin *et al.* 2005; Mosedale *et al.* 2019).

Multiallelic series comprised of multiple causal variants are more challenging. Our allele-based association approach only considers haplotype effects for a single genomic interval (i.e. the diplotype state probabilities do not vary in this region). Thus, it implicitly assumes that all causal variants are on the same genomic interval. For this reason, results from our allele-based approach cannot strictly be used to evaluate combinations of variants from different (even adjacent) genomic intervals.

An alternative approach assumes that the allelic series is at least as complicated as a given biallelic variant. In this case, the prior distribution of the allelic series is restricted to exclude partitions that violate the functional distinctions given by a causal variant. For example, for *J* = 3 haplotypes and a causal biallelic variant that contrasts haplotypes A and C with B, the prior distribution for the allelic series is

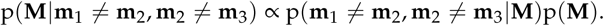

The first term on the right-hand side is an indicator variable denoting whether the allelic series **M** satisfies the conditions given by the biallelic variant, **m**_1_ ≠ **m**_2_ and **m**_2_ ≠ **m**_3_. The second term is the prior distribution of the allelic series (marginalized over the concentration parameter). Using this, we can compute a “variant-consistent” prior distribution that allows for multiallelic effects, but only in combination with the causal variant under consideration (and implicitly, only other variants on the same genomic interval, in proportion to the prior). This variant-consistent approach may be more useful than single-variant merge analysis for identifying candidate causal variants at multiallelic QTL. We implemented this variant-consistent prior distribution for the allelic series in our R package, and evaluating it would be an interesting topic for future research.

### Applying the Allele-Based Approach in Other Populations

Our allele-based association approach assumes that the underlying haplotypes at a QTL are known and that individuals can be probabilistically assigned to a diplotype (haplotype pair) state. These conditions are satisfied in diploid MPPs, where the founder haplotypes are known by construction, and our approach is designed for use in these populations. It is straight-forward to generalize our approach to haploid populations or polyploid MPPs. This simply involves defining the possible combinations of haplotypes (i.e. the number of columns in the **D** matrix) and an additive mapping of those combinations to haplotype frequency (the entries of a conformable **A** matrix). These generalizations can be implemented out of the box using our software.

In principle, our allele-based approach could be applied to any population, including human populations, provided that the underlying haplotypes at a QTL are known. In practice, haplotypes are typically unknown in non-experimental populations, and they must be defined empirically using combinations of adjacent variants (Meuwissen *et al.* 2014) or otherwise inferred as a reduced number of ancestral haplotypes (Davies *et al.* 2016; Pook *et al.* 2019). It is possible to define a fixed set of haplotypes and their probabilities *a priori* using another method and to provide this as input for our approach. Such a two-stage analysis would allow for allele-based inference in non-experimental populations, but it would not fully account for uncertainty in haplotype composition. It would also still be subject to computational constraints on the number of haplotypes and, in the case of the tree-informed prior, to caveats about local phylogeny in recombinant systems. Nonetheless, it would be interesting to apply our allele-based inference approach to QTL in non-experimental populations and to compare it with emerging haplotype-based, phylogeny-informed association approaches designed for these populations (Selle *et al.* 2020). This allelic perspective may provide new insight into the genetic architecture of QTL that is not revealed by the variant-based approaches commonly used in non-experimental populations.

### Limitations of the Allele-Based Approach

The allele-based association approach is limited by its computational speed. Our fully-Bayesian approach uses Gibbs sampling for posterior inference, which requires drawing many samples, at every locus, for every prior hypothesis. For the examples considered here, computation time for each analysis is on the order of minutes to hours. This limits the practical usefulness of the allele-based approach for applications such as QTL mapping. An alternative approach for the CRP may be approximate *maximum a posteriori* (MAP) inference (Raykov *et al.* 2016), which returns a single high-probability configuration of haplotypes. MAP avoids sampling and would be considerably faster than full posterior inference, though presumably with reduced performance.

Our method for calculating the tree-informed allelic series prior distribution is also computationally expensive. This is because it involves computing the prior probability of all 2^2*J*−2^ possible configurations of branch mutations **b** on a tree and recording the implied allelic series **M** for each. This approach is feasible for *J* = 8 founder haplotypes, the case for many MPPs, but not for *J* = 15, as in the DSPR. When *J* is large, it may be preferable to include the branch mutations **b** in the posterior sampling procedure, as in Azim Ansari and Didelot (2016), rather than integrating over them to precompute the prior distribution. We outline this approach in **Appendix C**. This requires mixing over the larger space of branch mutations **b**, though, rather than the smaller space of allelic series **M**, and the approach we outline only updates a single branch at a time. A full joint posterior sample of **b** is not tractable, but mixing could be improved by updating multiple branch mutations together, especially if sets of highly-dependent branches could be identified.

There are other possible alternatives for tree-informed allelic series inference. One such alternative is pseudo-marginal MCMC (Beaumont 2003; Andrieu and Roberts 2009). A pseudo-marginal approach would not sample the posterior branch mutations directly, but rather use a collection of branch mutation samples, weighed using importance sampling, to approximate the tree-informed allelic series prior distribution during posterior inference. Careful tuning of the proposal distributions in this framework could lead to efficient posterior sampling of the allelic series, but we have not explored this further. Another alternative is to disregard the explicit tree structure and instead use patristic distances between haplotypes as input for a distance dependent CRP (Blei and Frazier 2009), as in Cybis *et al*. (2018). It would be interesting to compare results from a distance dependent CRP with the tree-informed CRP that we have defined here. Yet another alternative is to identify a MAP set of branch mutations **b** or some other high-confidence set of mutated branches, as in Behr *et al*. (2020). This has the advantage of avoiding both computational bottlenecks (computing the tree-informed prior and sampling from the posterior), but the method described in Behr *et al*. (2020) does not account for the covariance between individuals induced by combinations of haplotypes and additive effects.

Lastly, the allele-based association approach only considers additive allele effects and unstructured error. As discussed in Jannink and Wu (2003), it is possible to include effects for allelic dominance in our model, though it would be desirable to include these as an additional variance component, as in Zhang *et al*. (2014). We did not consider error due to population structure in genetic background, which could also be included as an additional variance component (Eskin *et al.* 2008; Kang *et al*. 2010; Lippert *et al.* 2011; Zhou and Stephens 2012; Zhang *et al*. 2014). Adding this additional model complexity may be useful, but it would also increase the computational burden of the allele-based association approach.

## Acknowledgements

This work was primarily supported by the National Institute of General Medical Sciences under awards R01-GM104125 and R35-GM127000 (to W.V.) and T32-GM067553 (to W.L.C.) and the National Institute of Environmental Health Sciences under award R01-ES024965 (to S.N.P.K.). Computing resources were generously provided by the University of North Carolina Information Technology Services.

## Author contributions

W.L.C., S.N.P.K., and W.V. conceived the project and wrote the manuscript. W.L.C. wrote the software and performed the analyses. The authors declare no conflicts of interest.

## Appendix A: Including Covariates and Replicate Observations in the Likelihood

In this appendix, we generalize the likelihood function to account for optional covariates and replicate observations. The likelihood function is given by

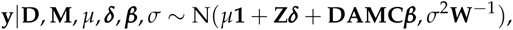

where **Z** is a *N* × *P* matrix of optional covariates, ***δ*** is a length *P* vector of covariate effects, **W** is a *N* × *N* diagonal matrix with the number of replicates for each observation on the diagonal, and the other variables are as previously described. We place a normal prior distribution on the covariate effects:

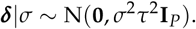

The likelihood can be rewritten as

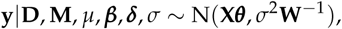

where **X** is the *N* × (*K* + *P*) design matrix, **X** = [**1 Z DAMC**], and ***θ***^*T*^ is a length *K* + *P* vector containing the intercept, independent effects, and covariate effects, ***θ***^*T*^ = [*µ* ***d***^*T*^ ***β***^*T*^]. The intercept, independent effects, and covariate effects are jointly distributed according to

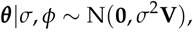

where **V** is a (*K* + *P*) × (*K* + *P*) diagonal matrix of the scaled prior covariance,

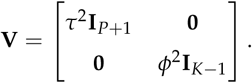

Conjugacy yields a closed form for a simplified, t-distributed likelihood function:

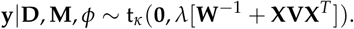

Posterior inference and prior elicitation are as previously described.

## Appendix B: Computing the Prior Distribution with the Tree-Informed CRP

In this appendix, we show that the prior density of the mutation status of the branches **b** can be marginalized over the concentration parameter *α* if the concentration parameter has a gamma prior distribution. This is useful for computing the tree-informed allelic series prior distribution. Our approach includes considerable bookkeeping of signs and coefficients, so we demonstrate using a minimal example, **b** = (0, 0, 1, 1, 1, 0) for *J* = 4.

We begin by expanding p(**b**|*T, α*):

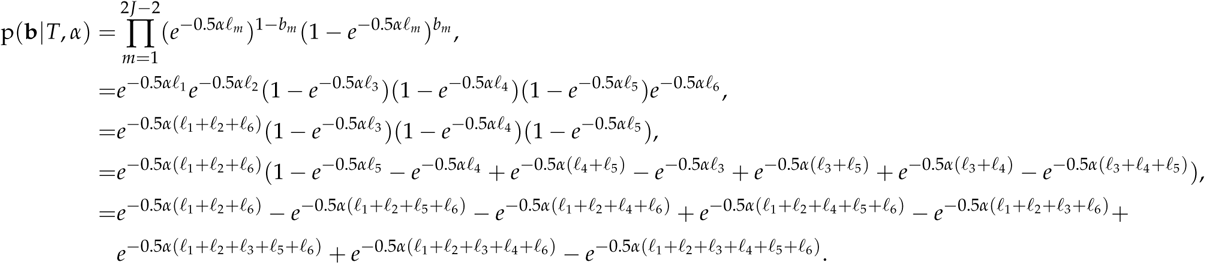

Each term is an exponentiated sum of branch lengths. All terms includes the branch lengths of the branches that are not mutated ({1, 2, 6}). The eight terms correspond to the eight possible subsets of the set of mutated branches ({2, 3, 4}), whose lengths are either included or excluded from the sum. If the sum includes an even number of mutated branch lengths, the sign of the term is positive, and if the sum includes an odd number, the sign is negative.

We are interested in

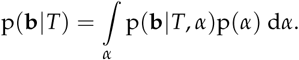

From the previous expansion, we know that this is an integral of a sum, which allows us to evaluate the integral separately for each term:

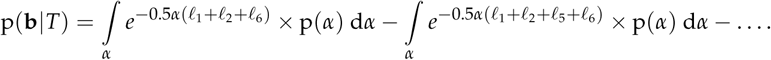

The prior distribution for the concentration parameter is

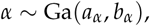

which has probability density function (omitting subscripts on the hyperparameters for clarity)

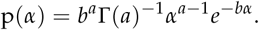

Focusing only on the first term of the expansion, we have

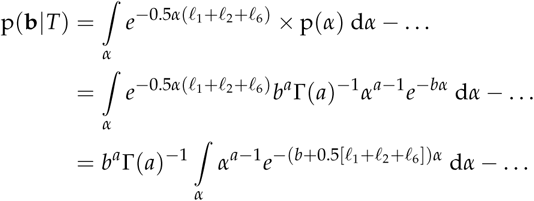

The quantity within the integral is the kernel of a gamma distribution with shape *a* and rate *b* + 0.5(*𝓁*_1_ + *𝓁*_2_ + *𝓁*_6_), and the integral is equal to the inverse of its normalizing constant:

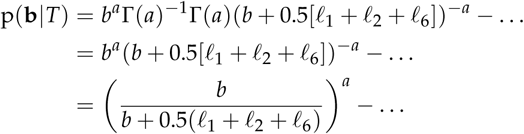

The other terms in the expression are solved similarly. This provides a closed form expression for p(**b**|*T*), which in turn is used to compute p(**M**|*T*).

## Appendix C: Posterior Sampling of Branch Mutations with the Tree-Informed CRP Prior Distribution

In this appendix, we outline an alternative approach for posterior inference of the allelic series, conditional on a tree *T*, that involves directly sampling the mutation status of the branches **b**. In the main text, we described an approach for posterior inference that has two steps: computing the marginal prior distribution of the allelic series **M** by integrating over all possible combinations of mutated branches (**Tree-Informed CRP**; **Appendix B**), and then conditionally updating the allele assignment of each haplotype, in proportional to the marginal prior distribution (**Posterior Inference 1b**). We use this approach because posterior inference is similar to the CRP, and we use the marginal prior distribution to calculate the marginal likelihood. However, computing the prior distribution is computationally prohibitive unless the number of founder haplotypes *J* is small, and this approach violates exchangeability during posterior inference. If either of these are a concern, it may be preferable to directly sample the mutation status of the branches during posterior inference. This avoids integrating over all possible combinations of mutated branches and does not violate exchangeability, but it does requires mixing over a larger state space, is conditional on a single tree, and does not facilitate calculating the marginal likelihood. We have implemented this approach but did not use it in our analyses.

The alternative approach for posterior inference of the allelic series is comprised of three steps: iteratively updating the mutation status of each branch, *b*_*m*_, conditional on the other branches, *b*_−*m*_; sampling the latent number of mutations on each branch, *γ*_*m*_, conditional on mutation status; and updating the mutation rate or concentration parameter, *α*, conditional on the total number of mutations. We now discuss each of these steps in detail.

### Updating the Mutation Status of Each Branch

First, we update the mutation status of each branch. The allelic series is completely determined by the tree and which branches are functionally mutated, **M** = *f* (*T*, **b**). Thus, the conditional posterior of the mutation status of a branch is given by

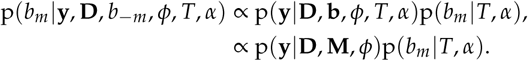

This is the product of the t-distributed likelihood and the categorical prior distribution of the mutation status of a branch. The conditional posterior is calculated directly for both settings of *b*_*m*_, and it reduces to the prior distribution of *b*_*m*_ when the allelic series is conditionally independent of the current branch, given the mutation status of the other branches: (**M *∐*** *b*_*m*_) | *b*_−*m*_.

### Sampling the Number of Mutations on Each Branch

Next, we sample the latent number of mutations on each branch. Functional mutations occur on the branches of the tree as a Poisson process with constant rate 0.5*α*. The prior number of mutations on each branch is a Poisson distribution with rate proportional to branch length:

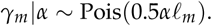

We used this relationship to construct the prior distribution of *b*_*m*_. By definition, if a branch is not functionally mutated, then there are no mutations on the branch,

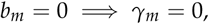

and if a branch is mutated, then the prior number of mutations on the branch is distributed as

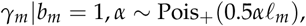

where Pois_+_ denotes the zero-truncated Poisson distribution. The conditional posterior number of mutations on all branches is

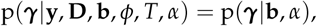

which is the joint conditional prior distribution and is straightforward to sample.

### Updating the Concentration Parameter

Finally, we update the concentration parameter (mutation rate), *α*, conditional on the total number of mutations. The conditional posterior for the concentration parameter is

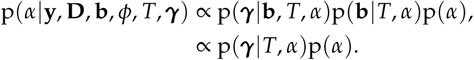

This is the product of a Poisson distribution for the number of functional mutations on the tree and the prior distribution for the concentration parameter. The posterior is gamma-distributed when the concentration parameter has a conjugate gamma prior distribution.

## Appendix D: Estimating the Marginal Likelihood

In this appendix, we describe an approach for estimating the marginal likelihood using the output of the Gibbs sampler (Chib 1995). Rearranging Bayes theorem, the log marginal likelihood can be expressed as

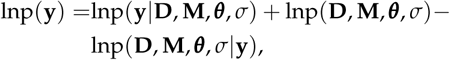

which is true at any point (**D**′, **M**′, ***θ***′, *σ*′). Optionally, these probabilities may also be conditional on a tree, *T*. Estimating the marginal likelihood involves factoring the joint posterior into terms that can be either well-approximated from the output of the Gibbs sampler or calculated directly.

### Estimating the Joint Posterior

An estimate of the joint posterior is given by

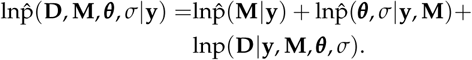

The first term, 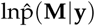, is the marginal posterior density of the allelic series. An approximation of the marginal posterior is given by the Gibbs sampler, and the density estimate at **M**′is the proportion of posterior samples equal to this value. This estimate is most accurate at the *maximum a posteriori* (MAP) allelic series, which we use to designate a point **M**′.

The second term of the estimate, 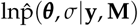, is the joint conditional posterior of the coefficients and the scale of the error variance. The joint posterior is conditional on the allelic series and must be evaluated at the point **M**′, but this distribution is not given directly by the Gibbs output. To obtain an estimate of the joint conditional posterior distribution, we resume iterating the Gibbs sampler after the initial posterior chain, but now with the allelic series fixed to **M**′. These conditional posterior samples yield sufficient statistics that we use to obtain an accurate Rao-Blackwellized estimate of the joint conditional posterior density (Blackwell 2007). We also use these sufficient statistics to designate ***θ***′ and *σ*′.

The final term of our estimate, lnp(**D** |**y, M, *θ***, *σ*), is the full conditional posterior of the diplotype states. This is calculated directly, and we designate the marginal MAP of the diplotype states for **D**′.

### Computing the Joint Prior

Factoring the joint prior is straightforward because most of the priors are independent by construction. The joint prior distribution is given by

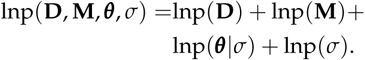

Calculating lnp(**M**) involves integrating over *α*. This must be approximated when using the CRP, but it is already calculated directly when conditioning on a tree. Calculating lnp(***θ***| *σ*) involves integrating over *ϕ*, which has a closed form involving a confluent hypergeometic function (Abramowitz and Stegun 1972). Finally, the last two terms in the prior are equal to zero when using an improper prior distribution for *µ* and *σ*, and they are only evaluated up to a constant. Despite this, BFs, which are ratios of marginal likelihoods, are still valid provided that these constants cancel (Servin and Stephens 2007).

### Supplemental Text: Alternative Prior Distributions for the Allelic Series

#### Alternatives Considered

In the main text, we focused on a specific prior distribution for the CRP concentration parameter, comparing it via simulation with the full haplotype-based association approach and evaluating the relative benefit of including tree information. In this supplement, we consider an alternative prior distribution for the concentration parameter, as well as an alternative prior distribution for the allelic series. The prior distributions considered in this supplement are:

- the one used in the main text, which is the CRP-based model with a relatively-conservative exponential prior distribution on the concentration parameter (termed the “Exponential” model here),

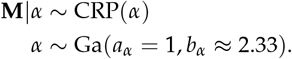

With *J* = 8 possible haplotypes, this prior distribution corresponds to a 50% probability of *K* = 1 functional allele, and monotonically favors smaller numbers of functional alleles *a priori*.
- the CRP-based model, but with a weakly informative gamma prior distribution on the concentration parameter (the “Gamma” model):

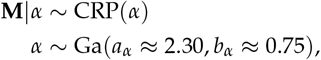

With *J* = 8 possible haplotypes, this prior distribution corresponds to a 5% probability of the null model with *K* = 1 functional allele and a 1% probability of the full model with *K* = 8 functional alleles.
- and a uniform prior which assumes that all configurations of the allelic-series are *a priori* equally likely (the “Uniform” model):

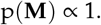

This is implemented as a non-exchangeable uniform process prior (Wallach *et al.* 2008).

These alternative prior distributions are shown in **Figure S1**. We evaluated these alternatives using the simulation procedure described in the main text.

#### Simulation Results

**Figure S2** shows the 0-1 accuracy of the MAP allelic series under the Uniform, Gamma and Exponential alternatives, for different numbers of true functional alleles and effect sizes. In the low power scenario (**A**), the ability to detect multiallelic series is low, but the two CRP-based approaches, Exponential and Gamma, have high accuracy when the QTL has only *K* = 1 functional allele (the null model), or when the QTL is biallelic. In the high power scenario (**B**), the CRP-based approaches have reasonably high accuracy (approximately 80%) for up to *K* = 4 functional alleles, with the Exponential outperforming the Gamma through this range. The more-diffuse Gamma prior maintains some limited accuracy through *K* = 8 alleles, outperforming the Exponential, although accuracy for highly-multiallelic series is generally low. In both scenarios, the Uniform approach is worse than the Exponential and Gamma, except when there is an intermediate number of alleles.

**Figure S3** is similar to the previous figure but shows the posterior probability of the correct allelic series, rather than the accuracy of the MAP allelic series. The Gamma and Uniform priors have relatively low certainty across both power scenarios and for all true numbers of functional alleles. In contrast, the Exponential prior is decisive when the true number of alleles is low, but at the expense of accuracy when the true number of alleles is high. Notably, in the high power scenario (**B**), the Gamma prior has reduced certainty when the QTL is null relative to biallelic, suggesting it has a tendency to overestimate the number of alleles under the null.

**Figure S4** is also similar to the previous figures but shows the posterior expectation of the number of alleles. In the low power scenario (**A**), the expectations for both the Gamma and Uniform are insensitive to the true number of alleles, consistently reporting an intermediate number of functional alleles. In contrast, the Exponential is accurate when the true number of alleles is one or two, but it reports approximately three alleles when the true number of alleles is higher. In the high power scenario (**B**), all the alternatives are more sensitive to the true number of alleles. In particular, the Exponential approaches the correct expectation for as many as *K* = 5 alleles. Notably, both the Exponential and Uniform show a tendency to overestimate when the QTL is null. We hypothesize that this is due to their relatively fat tails with respect to prior number of functional alleles, and that this acts in combination with a prior for QTL effect size that can accommodate small effects. When the true number of alleles is one, the QTL effect is necessarily zero, and it becomes “easier” for these permissive allelic series priors to estimate many effects, each of very small size.

**Figure S5** shows the MSE of haplotype effect estimates for the alternative allelic series prior distributions. In the low power scenario (**A**), the allele-based approaches outperform the Full haplotype-based approach when the true number of alleles is small. When power is high (**B**), this is also true when there are intermediate numbers of alleles. However, in both scenarios, when the true number of alleles is high, the Full model is better than the allele-based approach. As with accuracy, the Exponential is relatively better than Gamma and Uniform when the true number of alleles is low, and it is relatively worse when the true number of is high.

### Discussion

On the basis of these results, we recommend using the Exponential prior distribution for the allelic series. In many applied cases it will be reasonable to expect that QTL have only a few functional alleles, and when this is the case, the Exponential consistently outperforms than the other alternatives. In particular, when the number of alleles is small, the Exponential is less likely to overestimate the number of alleles than the Gamma or Uniform, and haplotype effect estimates are improved. Additionally, the Exponential prior typically provides a decisive posterior in the case of biallelic QTL. For these reasons, we focused on the Exponential prior distribution for the simulations in the main text.

### Supplemental Figures

**Figure S1.**
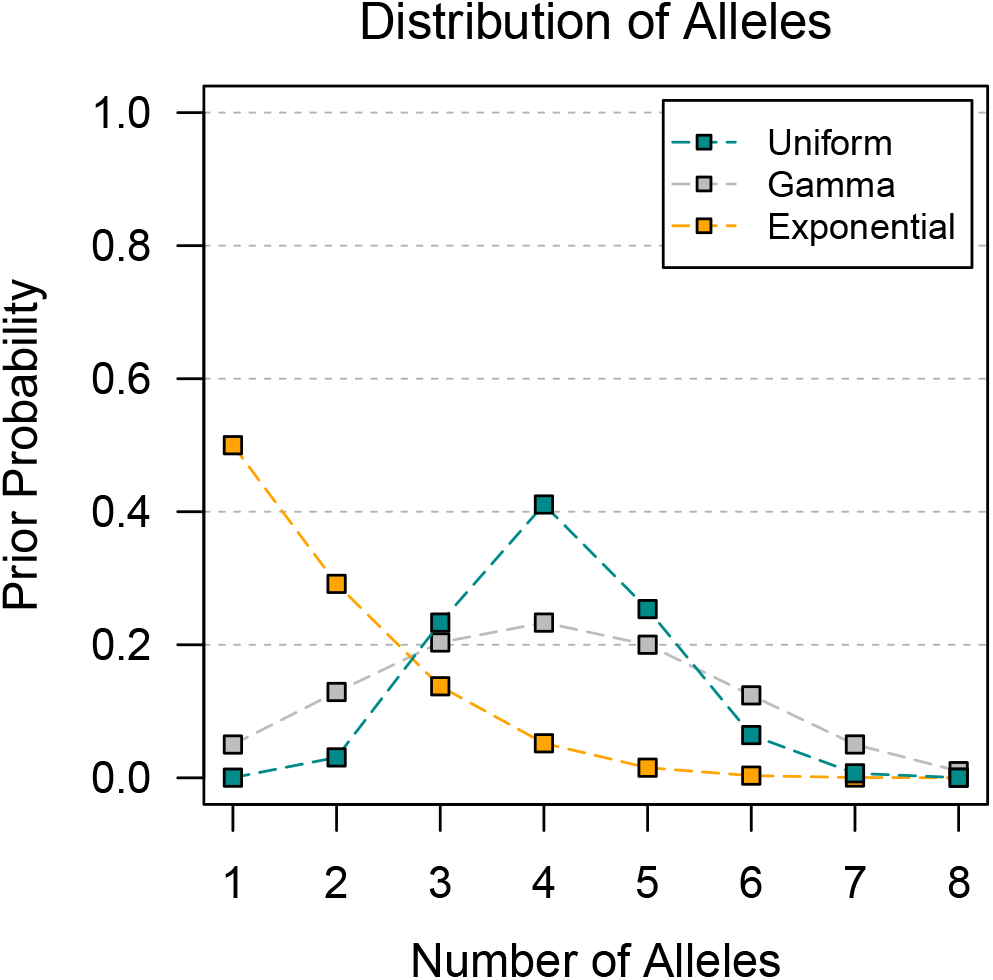
Prior distribution of number of functional alleles for the Uniform, Gamma, and Exponential prior distributions. Points are connected for clarity. Uniform places high prior weight on an intermediate number of functional alleles. Gamma is less informative than the Uniform with fatter tails. Exponential favors smaller numbers of alleles than the others.

**Figure S2.**
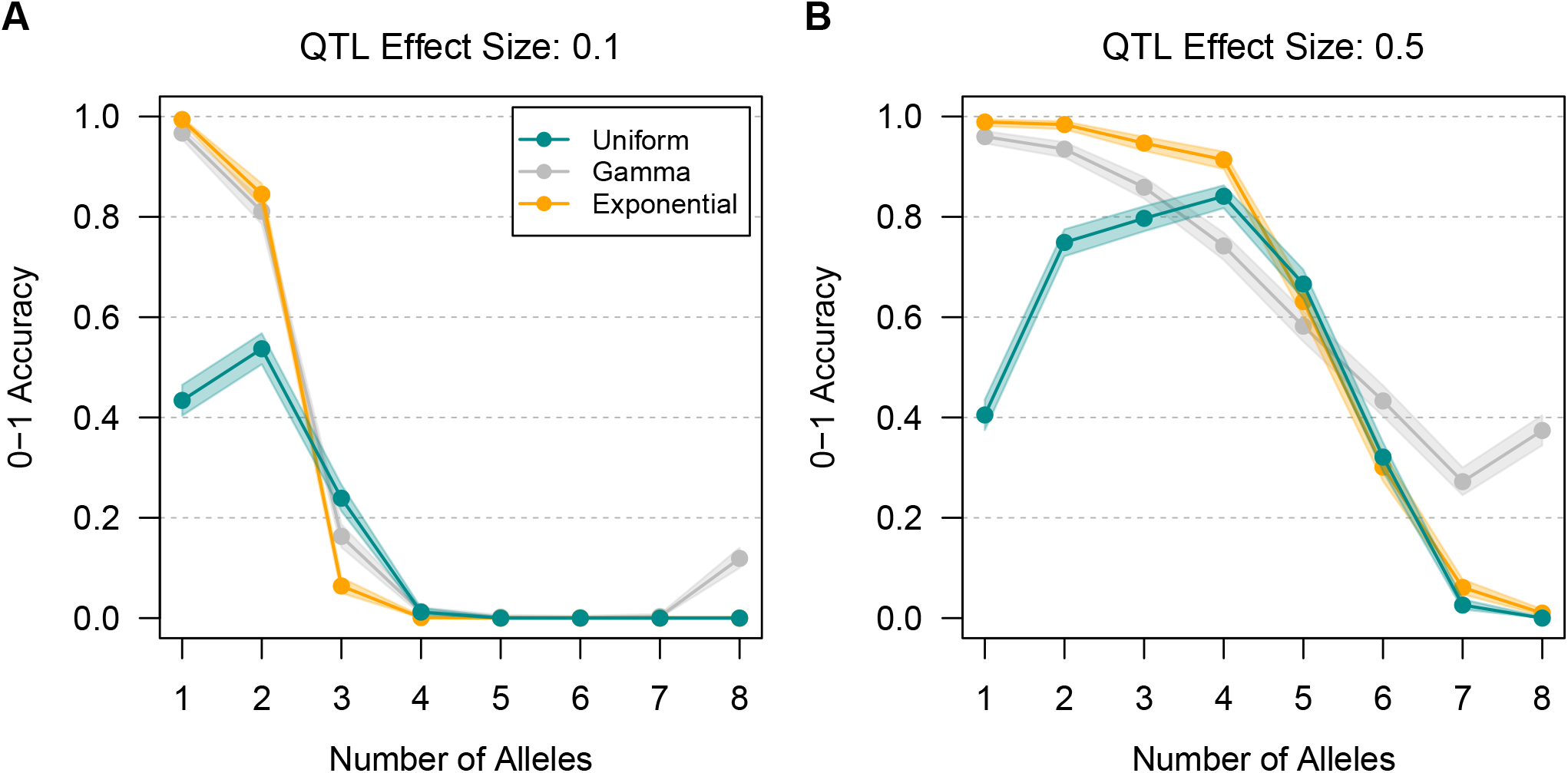
0-1 accuracy of posterior allelic series inference using the Uniform, Gamma, and Exponential prior distributions, for varying numbers of true functional alleles, across two effect sizes. Points are connected for clarity. Shading denotes 95% confidence intervals. In the low power scenario (**A**), accuracy for Exponential and Full is high when the QTL has two or fewer alleles but low when it is multiallelic. In the high power scenario (**B**), Exponential and Full have reasonable accuracy for an intermediate number of alleles, with the Exponential outperforming the Gamma through this range. Gamma maintains some accuracy for highly multiallelic series, outperforming the Exponential, although accuracy for highly-multiallelic series is generally low. Across both scenarios, Uniform is worse than Exponential and Gamma, except for an intermediate number of alleles.

**Figure S3.**
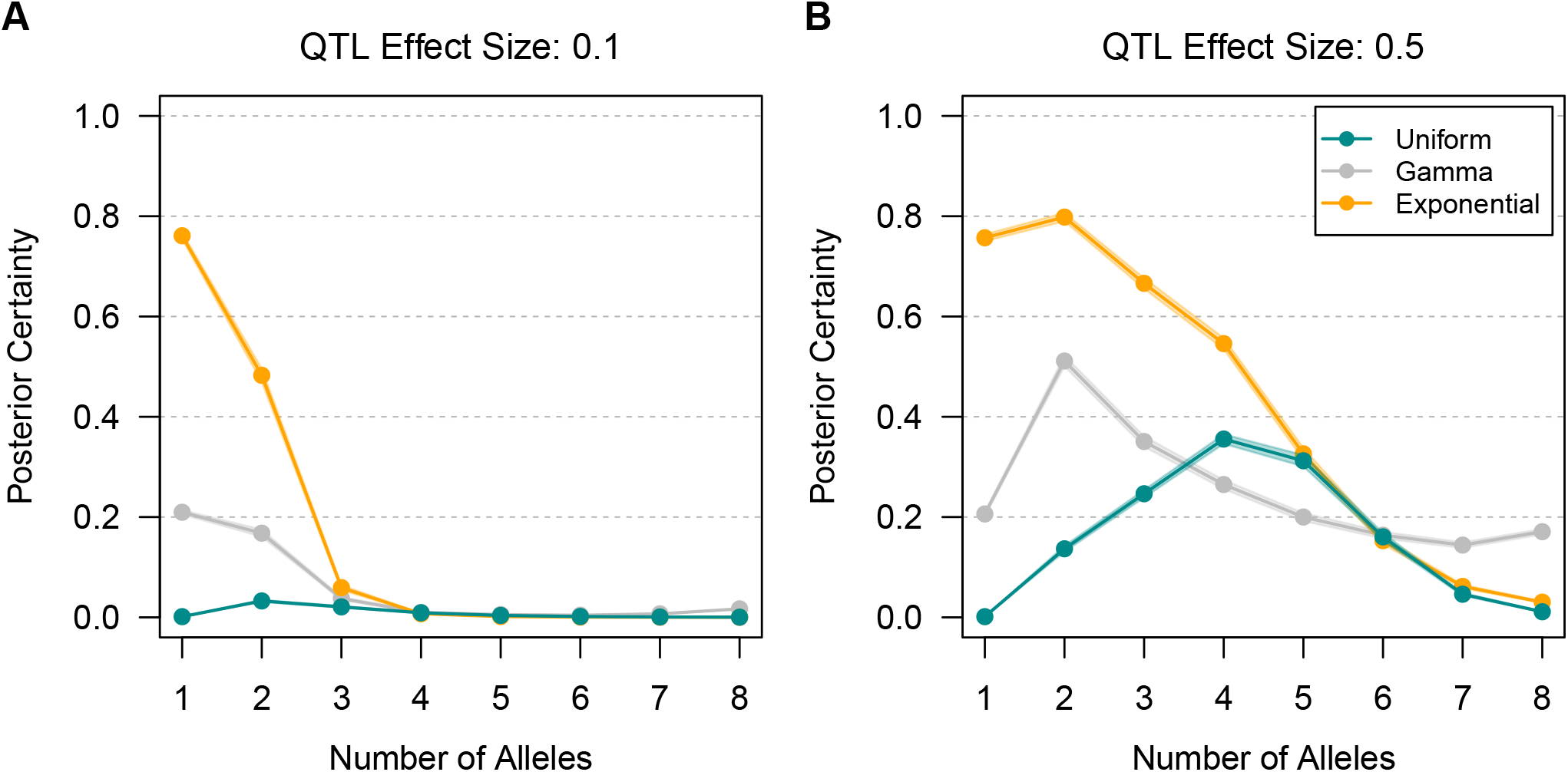
Posterior certainty of the correct allelic allelic series using the Uniform, Gamma, and Exponential prior distributions, for varying numbers of true functional alleles, across two effect sizes. Points are connected for clarity. Shading denotes 95% confidence intervals. In the low power scenario (**A**), posterior certainty for Exponential is high when the QTL has two or fewer alleles but low when it is multiallelic. Posterior certainty for Gamma and Uniform is low for any number of alleles. In the high power scenario (**B**), Exponential has higher posterior certainty for an intermediate number of alleles than Gamma and Uniform. Gamma has the highest posterior certainty when the QTL is highly multiallelic, but posterior certainty is low.

**Figure S4.**
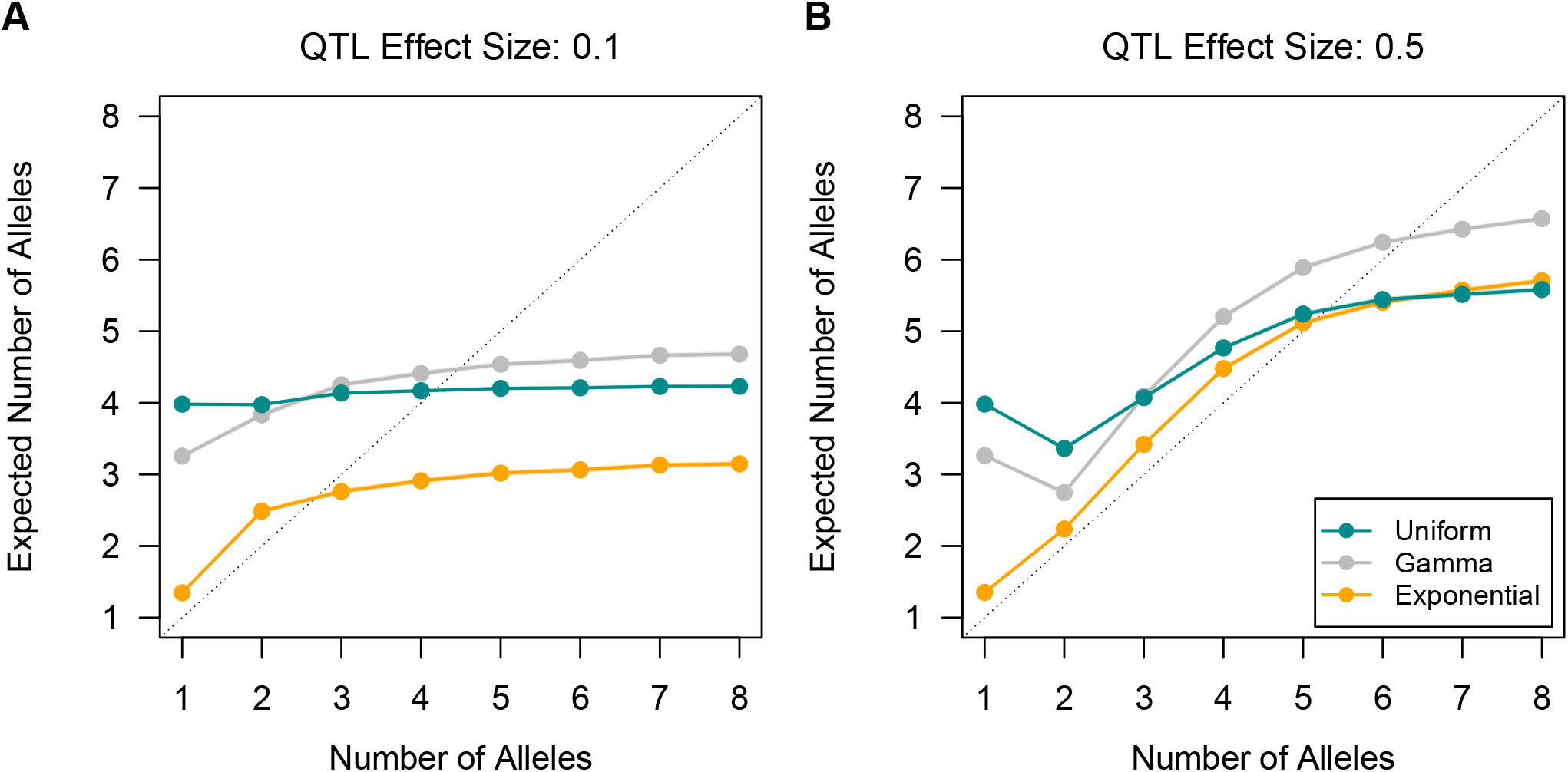
Posterior expectation of number of alleles using the Uniform, Gamma, and Exponential prior distributions, for varying numbers of true functional alleles, across two effect sizes. The dotted line indicates the correct expectation. Points are connected for clarity. Shading denotes 95% confidence intervals. In the low power scenario (**A**), Uniform and Gamma are relatively invariant and always expect an intermediate number of alleles. Exponential is better when the QTL has two or fewer alleles, but it underestimates the number of alleles for multiallelic series. In the high power scenario (**B**), Exponential is close to the correct expectation for an intermediate number of alleles but underestimates when the QTL is highly multiallelic. Gamma and Uniform are similar to Exponential but estimate more alleles, and they substantially overestimate when there is only one allele.

**Figure S5.**
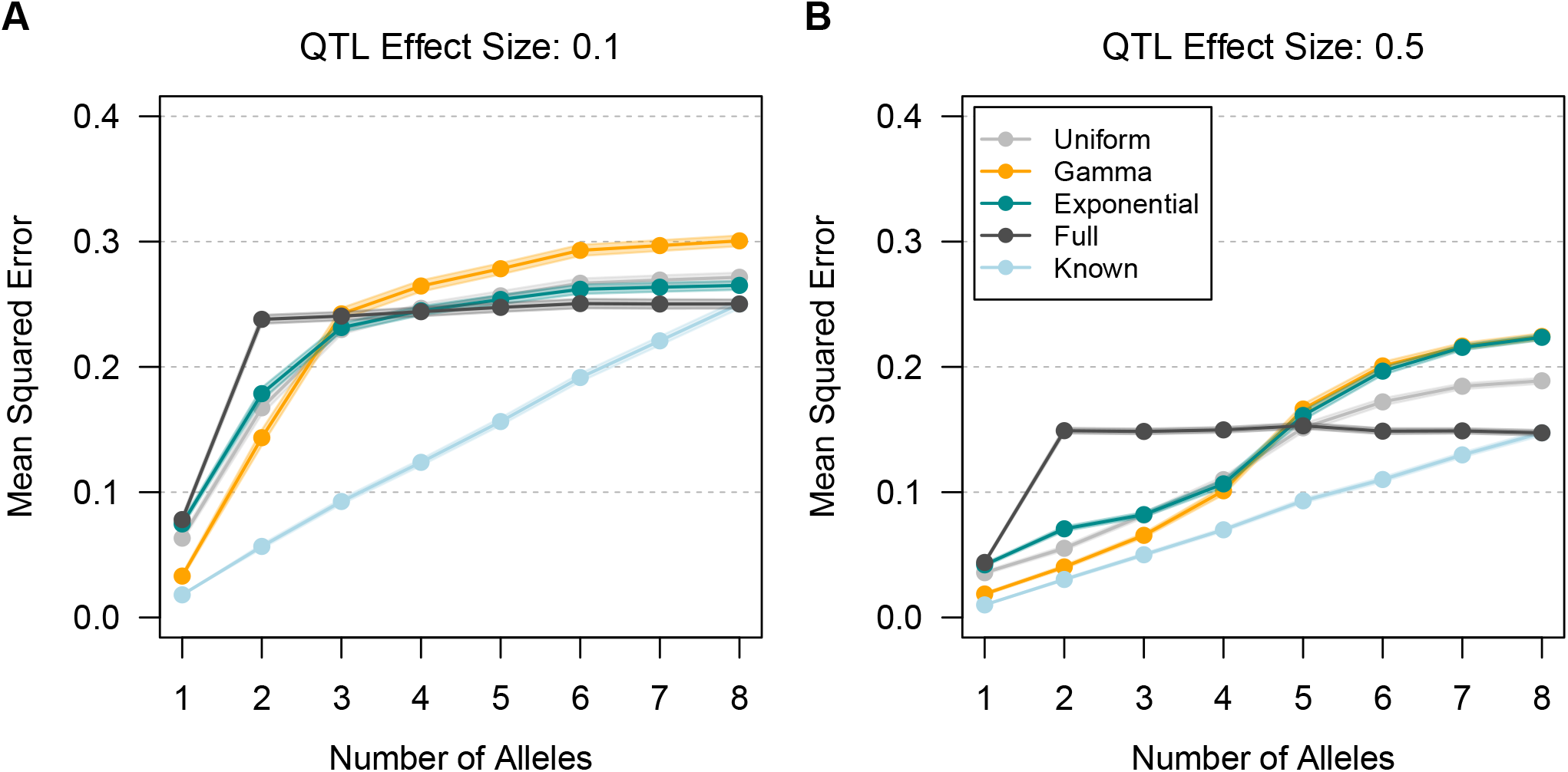
Mean squared error (MSE) of haplotype effect estimates using the Uniform, Gamma, and Exponential prior distributions, for varying numbers of true functional alleles, across two effect sizes. Full is the haplotype-based approach where all haplotypes are functionally distinct, and Known is an oracle prior in which the correct allelic series is known. Points are connected for clarity. Shading denotes 95% confidence intervals. In the low power scenario (**A**), Uniform, Gamma, and Exponential outperform Full when the true number of alleles is small. Exponential is better than Uniform and Gamma when the QTL has two or fewer alleles, but it is worse when the QTL is multiallelic. In the high power scenario (**B**), the Uniform, Gamma, and Exponential outperform Full for an intermediate number of alleles.

### Supplemental Tables

**Table S1.** Computation time in minutes for selected analyses.

| Analysis | # of Haplotypes | # of Samples | Duration (min) |
| --- | --- | --- | --- |
| PreCC MCV Full | 8 | 100,000 | 3.61 |
| PreCC MCV CRP | 8 | 100,000 | 8.01 |
| PreCC MCV Tree (prior) | 8 | (1000 trees) | 395.97 |
| PreCC MCV Tree (posterior) | 8 | 100,000 | 8.19 |
| DSPR CG4086 Full | 15 | 100,000 | 28.21 |
| DSPR CG4086 CRP | 15 | 1,000,000 | 410.99 |
| DSPR CG10245 Full | 15 | 100,000 | 29.67 |
| DSPR CG10245 CRP | 15 | 1,000,000 | 646.38 |

**Table S2.** Width of the 95% highest posterior density for MCV (fL) haplotype effects using the Full, CRP, and Tree approaches.

| Haplotype | Full | CRP | Tree |
| --- | --- | --- | --- |
| A | 6.60 | 4.06 | 2.99 |
| B | 5.03 | 3.36 | 3.23 |
| C | 7.14 | 5.04 | 3.04 |
| D | 5.87 | 3.44 | 3.23 |
| E | 5.96 | 3.43 | 3.23 |
| F | 5.96 | 5.57 | 5.72 |
| G | 6.08 | 3.96 | 4.43 |
| H | 5.07 | 3.74 | 2.99 |

**Table S3.**
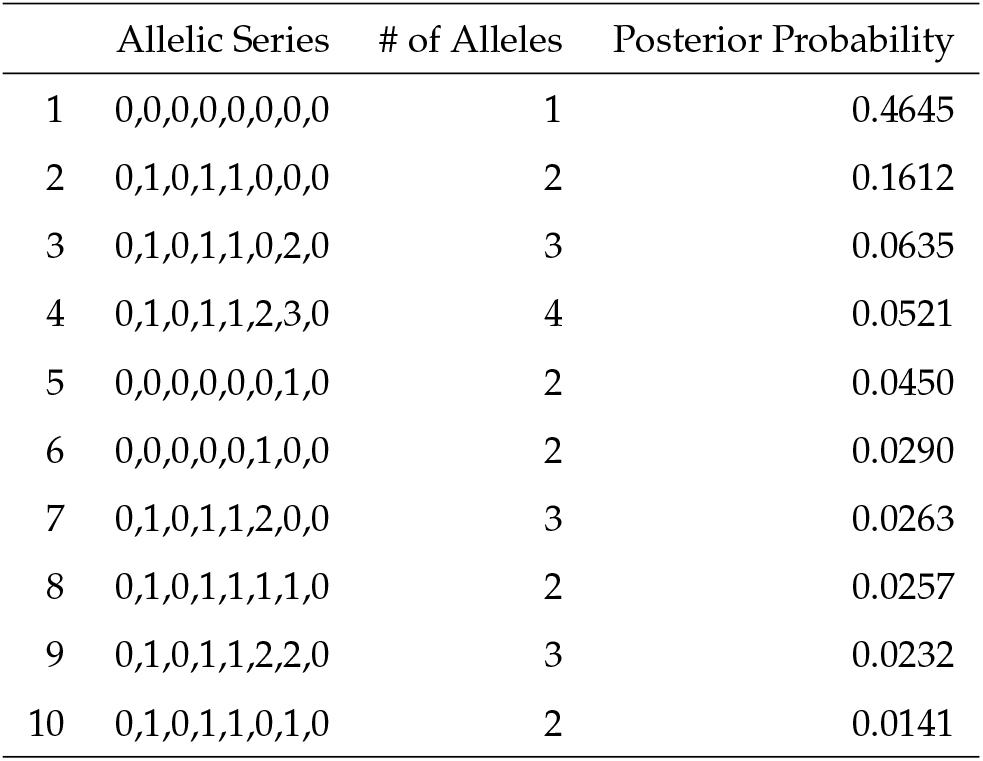
Top ten prior allelic series for MCV QTL in the PreCC using the Tree approach.

**Table S4.** Top ten posterior allelic series for CG4086 cis-eQTL in the DSPR using the CRP approach.

|  | Allelic Series | # of Alleles | Posterior Probability |
| --- | --- | --- | --- |
| 1 | 0,0,0,0,0,1,1,0,0,2,0,0,0,0,0 | 3 | 0.2536 |
| 2 | 0,0,0,0,0,1,2,0,0,3,0,0,0,0,0 | 4 | 0.0503 |
| 3 | 0,0,0,0,0,1,1,0,0,2,1,0,0,0,0 | 3 | 0.0482 |
| 4 | 0,0,0,0,1,1,1,0,0,2,0,0,0,0,0 | 3 | 0.0459 |
| 5 | 0,0,0,0,0,1,1,0,0,1,0,0,0,0,0 | 2 | 0.0361 |
| 6 | 0,1,1,1,1,0,0,1,1,2,1,1,1,1,1 | 3 | 0.0297 |
| 7 | 0,1,1,1,1,2,2,1,1,0,1,1,1,1,1 | 3 | 0.0253 |
| 8 | 0,0,0,0,0,1,1,0,0,2,2,0,0,0,0 | 3 | 0.0238 |
| 9 | 0,0,0,0,1,2,2,0,0,1,0,0,0,0,0 | 3 | 0.0229 |
| 10 | 0,0,0,0,1,1,1,0,0,2,1,0,0,0,0 | 3 | 0.0147 |

**Table S5.** Width of the 95% highest posterior density interval for CG4086 haplotype effects in the DSPR using the Full and CRP approaches.

| Haplotype | Full | CRP |
| --- | --- | --- |
| A1 | 3.56 | 2.99 |
| A2 | 1.19 | 0.17 |
| A3 | 1.03 | 0.17 |
| A4 | 0.99 | 0.17 |
| A5 | 5.11 | 3.11 |
| A6 | 1.81 | 1.31 |
| A7 | 1.07 | 0.68 |
| AB8 | 1.22 | 0.19 |
| B1 | 2.03 | 0.19 |
| B2 | 1.02 | 0.56 |
| B3 | 5.11 | 3.11 |
| B4 | 1.09 | 0.17 |
| B5 | 1.19 | 0.18 |
| B6 | 1.04 | 0.17 |
| B7 | 1.06 | 0.17 |

**Table S6.** Top ten posterior allelic series for CG10245 cis-eQTL in the DSPR using the CRP approach.

|  | Allelic Series | # of Alleles | Posterior Probability |
| --- | --- | --- | --- |
| 1 | 0,1,2,1,2,1,3,4,1,1,5,6,1,7,8 | 9 | 0.002380 |
| 2 | 0,1,2,3,2,1,3,3,3,3,4,1,3,0,2 | 5 | 0.001602 |
| 3 | 0,1,2,3,2,4,5,3,3,3,1,6,3,5,7 | 8 | 0.001206 |
| 4 | 0,1,2,3,2,1,4,3,3,3,1,5,3,6,0 | 7 | 0.001068 |
| 5 | 0,1,2,3,2,1,4,5,1,1,6,7,1,8,9 | 10 | 0.001064 |
| 6 | 0,1,2,1,2,1,3,4,4,1,1,5,4,6,0 | 7 | 0.001050 |
| 7 | 0,1,2,2,2,1,3,3,3,3,4,1,3,0,2 | 5 | 0.001040 |
| 8 | 0,1,2,3,2,4,5,3,3,3,1,6,3,7,5 | 8 | 0.001034 |
| 9 | 0,1,2,1,2,1,3,4,1,5,6,7,1,8,9 | 10 | 0.001030 |
| 10 | 0,1,2,2,2,1,3,4,1,1,5,6,1,7,8 | 9 | 0.001021 |

**Table S7.** Width of the 95% highest posterior density interval for CG10245 haplotype effects in the DSPR using the Full and CRP approaches.

| Haplotype | Full | CRP |
| --- | --- | --- |
| A1 | 1.10 | 1.56 |
| A2 | 2.42 | 2.61 |
| A3 | 1.33 | 1.83 |
| A4 | 9.95 | 7.76 |
| A5 | 1.11 | 1.72 |
| A6 | 1.01 | 1.59 |
| A7 | 0.88 | 1.53 |
| AB8 | 0.90 | 1.12 |
| B1 | 0.98 | 1.55 |
| B2 | 9.98 | 7.80 |
| B3 | 0.89 | 1.52 |
| B4 | 1.80 | 2.53 |
| B5 | 0.92 | 1.55 |
| B6 | 0.91 | 1.53 |
| B7 | 1.09 | 1.88 |

## Notes

https://github.com/wesleycrouse/TIMBR

### Competing Interest Statement

The authors have declared no competing interest.

## Literature Cited

Abramowitz, M. and I. Stegun, 1972 Handbook of Mathematical Functions with Formulas and Mathematical Tables. Courier Dover Publications.

Alberts, R., P. Terpstra, Y. Li, R. Breitling, J. P. Nap, et al., 2007 Sequence polymorphisms cause many false cis eQTLs. PLoS ONE 2.

Andrieu, C. and G. O. Roberts, 2009 The pseudo-marginal approach for efficient Monte Carlo computations. Annals of Statistics 37: 697–725.

Aylor, D. L., W. Valdar, W. Foulds-Mathes, R. J. Buus, R. a. Verdugo, et al., 2011 Genetic analysis of complex traits in the emerging Collaborative Cross. Genome Research 21: 1213–1222.

Azim Ansari, M. and X. Didelot, 2016 Bayesian inference of the evolution of a phenotype distribution on a phylogenetic tree. Genetics 204: 89–98.

Beaumont, M. A., 2003 Estimation of population growth or decline in genetically monitored populations. Genetics 164: 1139–1160.

Behr, M., M. Azim Ansari, A. Munk, and C. Holmes, 2020 Testing for dependence on tree structures. Proceedings of the National Academy of Sciences of the United States of America 117: 9787–9792.

Berestycki, N., 2009 Recent progress in coalescent theory. Ensaios Matematicos 16: 1–193.

Blackwell, D., 2007 Conditional Expectation and Unbiased Sequential Estimation. The Annals of Mathematical Statistics 18: 105–110.

Blei, D. M. and P. I. Frazier, 2009 Distance Dependent Chinese Restaurant Processes. Journal of Machine Learning Research 12: 2461–2488.

Bouckaert, R. and J. Heled, 2014 DensiTree 2: Seeing trees through the forest. bioRxiv pp. 1–11.

Broman, K. W., D. M. Gatti, P. Simecek, N. A. Furlotte, P. Prins, et al., 2019 R/qtl2: Software for Mapping Quantitative Trait Loci with High-Dimensional Data and Multiparent Populations. Genetics 211: 495–502.

Bult, C. J., J. A. Blake, C. L. Smith, J. A. Kadin, J. E. Richardson, et al., 2019 Mouse Genome Database (MGD) 2019. Nucleic Acids Research 47: D801–D806.

Cavanagh, C., M. Morell, I. Mackay, and W. Powell, 2008 From mutations to MAGIC: resources for gene discovery, validation and delivery in crop plants.

Chib, S., 1995 Marginal Likelihood from the Gibbs Output. Journal of the American Statistical Association 90: 1313.

Churchill, G. A., D. C. Airey, H. Allayee, J. M. Angel, A. D. Attie, et al., 2004 The Collaborative Cross, a community resource for the genetic analysis of complex traits. Nature Genetics 36: 1133–1137.

Collaborative Cross Consortium, F. A. Iraqi, M. Mahajne, Y. Salaymah, H. Sandovski, et al., 2012 The genome architecture of the Collaborative Cross mouse genetic reference population. Genetics 190: 389–401.

Crowley, J. J., Y. Kim, A. B. Lenarcic, C. R. Quackenbush, C. J. Barrick, et al., 2014 Genetics of adverse reactions to haloperidol in a mouse diallel: A drug-placebo experiment and Bayesian causal analysis. Genetics 196: 321–347.

Cybis, G. B., J. S. Sinsheimer, T. Bedford, A. Rambaut, P. Lemey, et al., 2018 Bayesian nonparametric clustering in phylogenetics: modeling antigenic evolution in influenza. Statistics in Medicine 37: 195–206.

Davies, R. W., J. Flint, S. Myers, and R. Mott, 2016 Rapid genotype imputation from sequence without reference panels. Nature Genetics 48: 965–969.

Degnan, J. H. and N. A. Rosenberg, 2009 Gene tree discordance, phylogenetic inference and the multispecies coalescent. Trends in Ecology and Evolution 24: 332–340.

Didion, J. P. and F. Pardo-Manuel de Villena, 2013 Deconstructing Mus gemischus: Advances in understanding ancestry, structure, and variation in the genome of the laboratory mouse. Mammalian Genome 24: 1–20.

Drummond, A. J., M. A. Suchard, D. Xie, and A. Rambaut, 2012 Bayesian phylogenetics with BEAUti and the BEAST 1.7. Molecular Biology and Evolution 29: 1969–1973.

Durrant, C. and R. Mott, 2010 Bayesian quantitative trait locus mapping using inferred haplotypes. Genetics 184: 839–852.

Escobar, M. D. and M. West, 1995 Bayesian density estimation and inference using mixtures. Journal of the American Statistical Association 90: 577–588.

Eskin, E., C. M. Wade, M. J. Daly, D. Heckerman, N. A. Zaitlen, et al., 2008 Efficient Control of Population Structure in Model Organism Association Mapping. Genetics 178: 1709–1723.

Ewens, W. J., 1972 The sampling theory of selectively neutral alleles. Theoretical Population Biology 3: 87–112.

Gatti, D. M., K. L. Svenson, A. Shabalin, L.-Y. Wu, W. Valdar, et al., 2014 Quantitative Trait Locus Mapping Methods for Diversity Outbred Mice. G3: Genes, Genomes, Genetics 4: 1623–1633.

Gelman, A., 2006 Prior distributions for variance parameters in hierarchical models. Bayesian Analysis 1: 515–533.

Haley, C. S. and S. A. Knott, 1992 A simple regression method for mapping quantitative trait loci in line crosses using flanking markers. Heredity 69: 315–324.

Hasegawa, M., H. Kishino, and T. aki Yano, 1985 Dating of the human-ape splitting by a molecular clock of mitochondrial DNA. Journal of Molecular Evolution 22: 160–174.

Huang, B. E., K. L. Verbyla, A. P. Verbyla, C. Raghavan, V. K. Singh, et al., 2015 MAGIC populations in crops: current status and future prospects. Theoretical and Applied Genetics 128: 999–1017.

Hudson, R. R. and N. L. Kaplan, 1985 Statistical properties of the number of recombination events in the history of a sample of DNA sequences. Genetics 111: 147–164.

Jannink, J.-L. and X.-L. Wu, 2003 Estimating allelic number and identity in state of QTLs in interconnected families. Genetical Research 81: 133–44.

Kamary, K., K. Mengersen, C. P. Robert, and J. Rousseau, 2014 Testing hypotheses via a mixture estimation model. 1412.2044.

Kang, H. M., J. H. Sul, S. K. Service, N. A. Zaitlen, S.-Y. Kong, et al., 2010 Variance component model to account for sample structure in genome-wide association studies. Nature Genetics 42: 348–354.

Kass, R. E. and A. E. Raftery, 1995 Bayes factors. Journal of the American Statistical Association 90: 773–795.

Keane, T. M., L. Goodstadt, P. Danecek, M. A. White, K. Wong, et al., 2011 Mouse genomic variation and its effect on phenotypes and gene regulation. Nature 477: 289–294.

Keele, G. R., W. L. Crouse, S. N. P. Kelada, and W. Valdar, 2019 Determinants of QTL Mapping Power in the Realized Collaborative Cross. G3: Genes, Genomes, Genetics 9: 1707–1727.

Kelada, D. S. N. P., M. D. E. Carpenter, D. D. L. Aylor, M. P. Chines, M. H. Rutledge, et al., 2014 Integrative Genetics of Allergic Inflammation in the Murine Lung. American Journal of Respiratory Cell and Molecular Biology 51: 436–445.

Kelada, S. N. P., D. L. Aylor, B. C. E. Peck, J. F. Ryan, U. Tavarez, et al., 2012 Genetic Analysis of Hematological Parameters in Incipient Lines of the Collaborative Cross. G3: Genes, Genomes, Genetics 2: 157–165.

Kelleher, J., Y. Wong, A. W. Wohns, C. Fadil, P. K. Albers, et al., 2019 Inferring whole-genome histories in large population datasets. Nature Genetics 51: 1330–1338.

King, E. G., S. J. Macdonald, and A. D. Long, 2012 Properties and power of the Drosophila synthetic population resource for the routine dissection of complex traits. Genetics 191: 935–949.

King, E. G., B. J. Sanderson, C. L. McNeil, A. D. Long, and S. J. Macdonald, 2014 Genetic dissection of the Drosophila melanogaster female head transcriptome reveals widespread allelic heterogeneity. PLoS Genetics 10: e1004322.

Kingman, J. F. C., 1982 On the genealogy of large populations. Journal of Applied Probability 19: 27–43.

Kingman, J. F. C., 2006 Random partitions in population genetics. Proceedings of the Royal Society of London. Series B. Biological Sciences 201: 217–217.

Kover, P. X., W. Valdar, J. Trakalo, N. Scarcelli, I. M. Ehrenreich, et al., 2009 A multiparent advanced generation inter-cross to fine-map quantitative traits in Arabidopsis thaliana. PLoS Genetics 5: e1000551.

Lilue, J., A. Shivalikanjli, D. J. Adams, and T. M. Keane, 2019 Mouse protein coding diversity: What’s left to discover? PLOS Genetics 15: e1008446.

Lippert, C., J. Listgarten, Y. Liu, C. M. Kadie, R. I. Davidson, et al., 2011 FaST linear mixed models for genome-wide association studies. Nature Methods 8: 833–837.

Macdonald, S. J. and A. D. Long, 2007 Joint estimates of quantitative trait locus effect and frequency using synthetic recombinant populations of Drosophila melanogaster. Genetics 176: 1261–1281.

Martínez, O. and R. N. Curnow, 1992 Estimating the locations and the sizes of the effects of quantitative trait loci using flanking markers. Theor. Appl. Genet. 85: 480–488.

Meuwissen, T. H., J. Odegard, I. Andersen-Ranberg, and E. Grindflek, 2014 On the distance of genetic relationships and the accuracy of genomic prediction in pig breeding. Genetics Selection Evolution 46.

Mosedale, M., Y. Cai, J. S. Eaddy, R. W. Corty, M. Nautiyal, et al., 2019 Identification of Candidate Risk Factor Genes for Human Idelalisib Toxicity Using a Collaborative Cross Approach. Toxicological Sciences 172: 265–278.

Mott, R., C. J. Talbot, M. G. Turri, A. C. Collins, and J. Flint, 2000 A method for fine mapping quantitative trait loci in outbred animal stocks. PNAS 97: 12649–12654.

Müller, P., F. A. Quintana, A. Jara, and T. Hanson, 2015 Bayesian Nonparametric Data Analysis. Springer Series in Statistics, Springer International Publishing, Cham.

Neal, R. M., 2000 Markov Chain Sampling Methods for Dirichlet Process Mixture Models. Journal of Computational and Graphical Statistics 9: 249–265.

Park, T. and D. A. Van Dyk, 2009 Partially collapsed Gibbs samplers: Illustrations and applications. Journal of Computational and Graphical Statistics 18: 283–305.

Pook, T., M. Schlather, G. de los Campos, M. Mayer, C. C. Schoen, et al., 2019 HaploBlocker: Creation of Subgroup-Specific Haplotype Blocks and Libraries. Genetics 212: 1045–1061.

Rasmussen, M. D., M. J. Hubisz, I. Gronau, and A. Siepel, 2014 Genome-Wide Inference of Ancestral Recombination Graphs. PLoS Genetics 10: e1004342.

Raykov, Y. P., A. Boukouvalas, and M. A. Little, 2016 Simple approximate MAP inference for Dirichlet processes mixtures. Electronic Journal of Statistics 10: 3548–3578.

Robert, C. P., 2016 The expected demise of the Bayes factor. Journal of Mathematical Psychology 72: 33–37.

Rota, G.-C., 1964 The Number of Partitions of a Set. The American Mathematical Monthly 71: 498–504.

Selle, M. L., I. Steinsland, F. Lindgren, V. Brajkovic, V. CubricCurik, et al., 2020 Hierarchical modeling of haplotype effects based on a phylogeny. bioRxiv p. 2020.01.31.928390.

Servin, B. and M. Stephens, 2007 Imputation-based analysis of association studies: Candidate regions and quantitative traits. PLoS Genetics 3: 1296–1308.

Thompson, K. L. and L. S. Kubatko, 2013 Using ancestral information to detect and localize quantitative trait loci in genome-wide association studies. BMC Bioinformatics 14: 200.

Valdar, W., C. C. Holmes, R. Mott, and J. Flint, 2009 Mapping in structured populations by resample model averaging. Genetics 182: 1263–1277.

Valdar, W., L. C. Solberg, D. Gauguier, S. Burnett, P. Klenerman, et al., 2006 Genome-wide genetic association of complex traits in heterogeneous stock mice. Nature Genetics 38: 879–887.

van Dyk, D. A. and T. Park, 2008 Partially Collapsed Gibbs Samplers: Theory and Methods. Journal of the American Statistical Association 103: 790–796.

Wallach, H. M., S. T. Jensen, L. Dicker, and K. A. Heller, 2008 An Alternative Prior Process for Nonparametric Bayesian Clustering. Proceedings of the Thirteenth International Conference on Artificial Intelligence and Statistics (AISTATS) pp. 892–899.

Wei, J. and S. Xu, 2016 A Random-Model Approach to QTL Mapping in Multiparent Advanced Generation Intercross (MAGIC) Populations. Genetics 202: 471–86.

Welling, M., 2006 Flexible Priors for Infinite Mixture Models. In Proceedings of the Workshop on Learning with Nonparametric Bayesian Methods, 23rd ICML.

Williams IV, R., J. E. Lim, B. Harr, C. Wing, R. Walters, et al., 2009 A common and unstable copy number variant is associated with differences in Glo1 expression and anxiety-like behavior. PLoS ONE 4.

Yalcin, B., J. Flint, and R. Mott, 2005 Using progenitor strain information to identify quantitative trait nucleotides in outbred mice. Genetics 171: 673–681.

Yang, H., J. R. Wang, J. P. Didion, R. J. Buus, T. A. Bell, et al., 2011 Subspecific origin and haplotype diversity in the laboratory mouse. Nature Genetics 43: 648–655.

Yuan, Z., F. Zou, and Y. Liu, 2011 Bayesian multiple quantitative trait loci mapping for recombinantinbred intercrosses. Genetics 188: 189–195.

Zhang, Z., W. Wang, and W. Valdar, 2014 Bayesian Modeling of Haplotype Effects in Multiparent Populations. Genetics 198: 139–156.

Zhang, Z., X. Zhang, and W. Wang, 2012 HTreeQA: Using Semi-Perfect Phylogeny Trees in Quantitative Trait Loci Study on Genotype Data. G3: Genes, Genomes, Genetics 2: 175–189.

Zheng, C., M. P. Boer, and F. A. van Eeuwijk, 2015 Reconstruction of Genome Ancestry Blocks in Multiparental Populations. Genetics 200: 1073–1087.

Zhou, X. and M. Stephens, 2012 Genome-wide efficient mixed-model analysis for association studies. Nature Genetics 44: 821–824.

